# Developmental plasticity facilitates the structural maturation of cochlear inner hair cell ribbon synapses

**DOI:** 10.1101/2025.08.19.671016

**Authors:** Roos A. Voorn, Michael Sternbach, Jerome Bourien, Noboru H. Komiyama, Vladan Rankovič, Fred Wolf, Seth G. N. Grant, Christian Vogl

**Affiliations:** Auditory Neuroscience Group, Institute of Physiology, Medical University Innsbruck, 6020 Innsbruck, Austria; Presynaptogenesis and Intracellular Transport in Hair Cells Junior Research Group, Institute for Auditory Neuroscience and InnerEarLab, University Medical Center Goettingen, 37075 Goettingen, Germany; Collaborative Research Center 889 “Cellular Mechanisms of Sensory Processing”, 37075 Goettingen, Germany; Göttingen Graduate Center for Neurosciences, Biophysics and Molecular Biosciences, 37075 Goettingen, Germany; Max Planck Institute for Dynamics and Self-Organization, 37077 Goettingen, Germany; Göttingen Campus Institute for Dynamics of Biological Networks, University of Göttingen, 37073 Goettingen, Germany; Bernstein Center for Computational Neuroscience, 37073 Goettingen, Germany; INM, University of Montpellier, INSERM, Montpellier, France; Genes to Cognition Program, Centre for Clinical Brain Sciences, University of Edinburgh, Edinburgh EH16 4SB, UK; Simons Initiative for the Developing Brain, Centre for Discovery Brain Sciences, University of Edinburgh, Edinburgh EH8 9XD, UK; The Patrick Wild Centre for Research into Autism, Fragile X Syndrome & Intellectual Disabilities, Centre for Discovery Brain Sciences, University of Edinburgh, Edinburgh EH8 9XD, UK; Muir Maxwell Epilepsy Centre, University of Edinburgh, Edinburgh EH8 9XD, UK; Institute for Auditory Neuroscience and InnerEarLab, University Medical Center Goettingen, 37075 Goettingen, Germany; Otto Creutzfeldt Fellow of the Elisabeth and Helmuth Uhl Foundation

**Author notes:** Correspondence should be addressed to: Christian Vogl. UCB Pharmaceuticals, 1070 Brussels, Belgium.

**Keywords:** Cochlear hair cells, synaptogenesis, sensory systems, maturational refinement

## Abstract

Sound detection occurs in the cochlea, where sensory inner hair cells (IHC) accurately convert auditory stimuli into neurochemical signals. Presynaptically, IHCs harbor synaptic ribbons, specialized scaffolds that facilitate ultrafast and indefatigable exocytosis. During synapse assembly and subsequent maturation, IHC ribbons increase in volume and synaptic vesicle tethering capacity. This development is thought to result from progressive precursor aggregation. However, the underlying mechanisms of ribbon synapse formation have remained elusive thus far. In this study, we established a novel triple-color live-cell imaging approach to monitor IHC pre-synaptogenesis *in situ*. We found that ribbon precursors are highly dynamic and undergo bidirectional plasticity. The presynaptic active zone (AZ) forms a focal point for dramatic structural remodeling of precursors, which the AZ recruits, confines and redistributes. Furthermore, silencing spontaneous synaptic activity decreased precursor mobility and plasticity at the AZ. This suggests a fundamental role for activity-dependent Ca^2+^ influx in the plastic development shaping the unique properties of auditory ribbon synapses.

## Introduction

Ultrafast synaptic sound encoding translates physical sound waves into neural code, thus enabling speech perception and spatial orientation in a three-dimensional environment. In mammals, this challenging task is performed by ‘ribbon-type’ synapses between cochlear inner hair cells (IHC) and postsynaptic spiral ganglion neurons (SGN) that ensure temporally-precise and indefatigable signal transduction. Ribbons play a fundamental role in the recruitment, priming, and replenishment of synaptic vesicles (SVs), and may further serve as a diffusion barrier for presynaptic Ca^2+^ at the active zone (AZ) ^1–8^. In fact, ribbons are thought to fulfill a multifaceted scaffolding role at the cytomatrix of the AZ (CAZ), where they tether glutamate-filled SVs and cluster presynaptic Ca_V_1.3 channels – thus facilitating ultrafast and highly efficient excitation-secretion coupling ^9,10^.

While various studies have investigated the unique structure and physiology of mature IHC ribbon synapses, knowledge on the early development remains limited to date. Yet, previous findings indicate a dynamic process in which small cytoplasmic ribbon precursors gradually accumulate at the assembling AZ during late embryogenesis and gradually fuse to form mostly single, membrane-anchored ribbons at hearing onset ^11–13^. This structural transition period is currently thought to involve molecular motor driven re-distribution of ribbon material that largely concludes around P12 ^13,14^. However, the exact underlying molecular mechanisms as well as the regulatory pathways remain to be determined. Notably, it is well established that developing organs of Corti intrinsically generate spontaneous activity (SA) waves, which trigger sensory-independent IHC exocytosis to initiate auditory pathway activation. While this pre-sensory SA has been proven essential for the diversification and survival of SGNs ^15–17^, it remains unclear to which degree this process regulates the maturational refinement of the presynaptic architecture of IHC ribbon synapses. Yet, activity-induced adaptations in ribbon structure have previously been reported in other sensory systems ^18–22^.

To now experimentally address these issues, we set out to investigate IHC ribbon synapse development in real time during the critical early postnatal period. For this purpose, we devised a multi-color live-cell imaging paradigm to monitor the structural dynamics of IHC pre- and postsynaptic components *in situ* using short-term organotypically-cultured cochlear explants as a model system and combined this with Ca^2+^ imaging, immunohistochemistry and optical nanoscopy to comprehensively characterize developmental maturation of IHC AZs. We recently found that auditory ribbon synapse assembly is a highly dynamic process, in which ribbon precursors are subject to regular bidirectional plasticity events – i.e., precursor fusion and fission 14. Here, we show that the presynaptic AZ serves as a plasticity hub, where ribbon precursors can be recruited and stabilized, yet also locally re-distributed or even exchanged between distant AZs in a strikingly targeted manner. Moreover, the observed structural plasticity appears to depend on synapse location within the IHC basolateral compartment and is further regulated by SA-driven presynaptic Ca^2+^ influx.

## Materials & Methods

### Animals

All animal experiments were conducted in accordance with national as well as regional guidelines of Lower Saxony, Germany and were approval by the Lower Saxony animal welfare officer. For our experiments, PSD95-HaloTag23,24 mice were crossbred with Ai32-Vglut3-Cre Knock-In mice (Ai32-VC-KI-PSD ^14,25–27;^ Ai32: JAX#024109), or Ai14-Neurogenin1-Cre/ERT2 mice (Ai14-Neurog1-PSD ^25,28^; Ai14: JAX#007914; Neurogenin1-Cre: JAX#008529). Furthermore, C57Bl6/J wild-type mice (WT), and SNAP25-2A-GCaMP6s-D mice (SNAP25-GCaMP6 ^29^; SNAP25: JAX#025111) were used. Mice of either sex between the ages of postnatal day (P)3 and 7 were sacrificed by decapitation for preparation of organotypic cultures of the organ of Corti.

### Organotypic culture preparation of the mouse organ of Corti

Organotypic cultures of the organ of Corti were prepared from neonatal mice (P3-P7), of WT, Ai14-Neurog1-PSD, Ai32-VC-KI-PSD, and SNAP25-GCaMP6 background ^14^. Apical-medial turns of the organ of Corti were dissected out of the mouse cochlea and mounted on either 1.5 thickness high-precision coverslips (within a 35 mm Petri dish) or glass bottom Petri dish inserts (P35G-1.5-14-C, MatTek), coated with CellTak (#354240, Corning, 1:8 solution in NaHCO3). Upon proper attachment, the organotypic cultures were submerged in 2 mL culturing medium and incubated at 37°C, 5% CO_2_ for up to three days *in vitro* (DIV3).

### Viral transduction

Viral transduction *in vivo* was conducted by injection of neonatal animals of the *Ai14-Neurog1-PSD* and *Ai32-VC-KI-PSD* background with an adeno-associated virus vectors (AAV) PHP.B/eB (full constructs consisting of: PHP.B_CMV-HBA-RIBEYE-GFP_WRPE_bGH or PHP.eB_CMV-HBA_RIBEYE-tdTomato_WPRE_ bGH), one day prior to culture preparation (P4-P6) as described previously 14,30. Viral injections were performed through the round window of the right ear under 5% isoflurane anesthesia (5% for induction, 2-3% for maintenance). Analgesia consisted of subcutaneous injection of buprenorphine (0.1 mg/kg body weight, approximately 30 minutes before the surgery), locally applied xylocain (10 mg spray, during surgery), and subcutaneous injections of carprofen (5 mg/kg body weight, applied during as well as 1-day post-surgery). All virus injection experiments were performed in compliance with the national animal care guidelines and were approved by both the board for animal welfare of the University Medical Center Göttingen and the State Animal Welfare office of Lower Saxony (AZ19/3133).

### Application of JaneliaFluor HaloTag ligands and pharmacological treatment

Fluorescent live-cell labeling of the postsynaptic density (PSD) was conducted in our PSD95-HaloTag knock-in mouse lines using either the JaneliaFluor HaloTag JF549 (JF549, #GA111A, Janelia Fluor® HaloTag®) or the JaneliaFluor HaloTag 646 ligands (JF646, #GA112A, Janelia Fluor® HaloTag®) – final concentration: 200 nM. The JF549 ligand was used in dual-color experiments in combination with overexpression of RIBEYE-GFP; the JF646 ligand was used in the triple-color experimental setup, either combined with either RIBEYE-GFP or RIBEYE-tdTomato. The organotypic cultures were incubated with the ligand for 1.5 hours, after which the complete media was replaced in one washing step and the sample was incubated for 3-8 hours (37°C; 5% CO_2_) to wash out excess fluorophore, prior to imaging.

Pharmacological treatment with the CaV antagonist isradipine (#2004, Tocris) was performed by drop-wise bulk application of 100 μL culture media to a final concentration of 10 μM, in between time-lapse imaging sessions.

### Long-term multi-color time-lapse imaging

Time-lapse live-cell imaging experiments were conducted at a Nikon Eclipse Ti-E microscope equipped with a Yokogawa CSU-WI Spinning Disk setup and Andor laser system, under environmentally-controlled conditions (37°C, 5% CO_2_, Okolab Bold Line Cage Incubator), at 60x water immersion, NA 1.20. To prevent evaporation during long-term imaging, we used RI-matched silicon oil instead of water for objective immersion (Immersol W 2010, Ne 1.33). Three-dimensional regions of interest (ROI) were selected for high intensity PSD-HaloTag labeling, low to moderate RIBEYE-tdTomato or RIBEYE-GFP expression, and, when present, genetic fluorescence indicating healthy morphology of IHCs or SGNs. ROI dimensions were minimized, while including a margin for potential drift. This resulted in an axial range of approximately 29 μm and varying spatial dimension (15-30 μm x and/or y), adapted to tissue orientation. Multi-color 3D time-lapse imaging was performed at maximum acquisition speed (consecutive color acquisition), conducting continuous imaging for a period of 40-90 minutes. For dual-color experiments, time-lapse images were acquired at intervals of 45-60 seconds, whereas triple-color acquisition required 70-80 seconds per time frame. For dual-color experiments, laser excitation was used at 488 and 565 nm (20ms and 40ms). For triple-color experiments, excitation was dependent on the experimental setup and mouse line used: for experiments with the Ai14-Neurog1-PSD line, excitation of 488, 565 and 637 nm was used; in case of the Ai32-VC-KI-PSD line, excitation was used at 515, 565 and 637 nm (20ms, 20ms and 40ms respectively, in both experimental setups). All imaging frames were averaged 4 times.

### Immunohistochemistry of organotypically-cultured or acutely-dissected organs of Corti

Organotypic cultures were fixed in 4% formaldehyde (in PBS) for 15 minutes on ice, and thereafter stored in PBS at 4°C. Fixed organs of Corti were permeabilized in PBS + 0.5% TritonX-100 for 30 minutes, then incubated in blocking buffer (PBS + 0.5% Triton-X100 + 10% normal goat serum) for 1 hour at room temperature. Incubation with primary as well as secondary, or directly-conjugated antibodies was performed in blocking buffer over 2 hours at room temperature, while protected from light. Used antibodies can be found in the supplementary information (Table ST1). Live-cell imaging samples on glass bottom dishes were enclosed with an additional coverslip, whereas samples on coverslips were mounted directly on glass slides, in both cases using ProLong Glass Antifade reagent (#P36984, Invitrogen).

### Confocal and super-resolution STED microscopy of fixed samples

Immunostained samples were imaged using an Abberior Instruments Expert Line STED microscope – operated in confocal laser scanning mode, using either a 20x/NA 0.8 or 100x/NA 1.4 oil immersion objective. For super-resolution STED microscopy, samples were imaged in 2D STED mode at a 10 nm xy pixel size. A pulsed 775 nm laser was used for stimulated emission depletion, while using 561 nm and 640 nm laser lines for excitation.

### Processing and analysis of confocal and STED images of fixed samples

Confocal images of fixed organs of Corti were analyzed for ribbon volume and postsynaptic attachment in IMARIS (9.6.1, Oxford Instruments), using the Surface rendering function for ribbons (0.054 surface detail, local background subtraction, 0.28 μm largest sphere diameter, 0.15 split surface seed points, 3 quality filter), and for the PSD (0.08 surface detail, local background subtraction, 0.6 μm largest sphere diameter, 3 quality filter). Based on their proximity to the PSD, ribbon precursors were either classified as synaptically-engaged (≤500 nm), or non-synaptic (>500 nm). Super-resolution STED images were processed for display in FIJI/ImageJ (2.3.0/1.53q) using the Gaussian Blur filter at σ 1.0.

### Time-lapse image processing and analysis

Time-lapse images were post-processed in FIJI/ImageJ (2.3.0/1.53q) to correct for bleaching using the BleachCorrection plugin, simple ratio algorithm (CorrectBleach ^31^); different fluorophores were corrected separately. Thereafter, two consecutive drift corrections were performed in IMARIS (9.6.1, Oxford instruments); the first correction based on the surface rendering of the cellular context (IHC, SGN, or PSD), the second correction originating from ribbon precursor particle tracking using the Spots and lineage tracing algorithm.

The following surface rendering parameters were used: for SGNs (0.22 surface detail, local background subtraction, 0.5 μm largest sphere diameter, 10 quality filter, autoregression tracing, 1.5 μm maximum distance, 0 maximum time gap), IHCs (0.26 surface detail, local background subtraction, 0.2 μm largest sphere diameter, 10 quality filter, autoregression tracing, 1.5 μm maximum distance, 0 maximum time gap) and PSDs (0.16 surface detail, 165 local background subtraction, 0.5 μm largest sphere diameter, 10 quality filter, autoregression tracing, 1.5 μm maximum distance, 0 maximum time gap). When the PSD-HaloTag labeling was used in combination with genetic YFP IHC plasma membrane fluorescence (Ai32-VC-KI-PSD), the PSD surface rendering was filtered for proximity to the IHC surface; when the PSD-HaloTag label was used in combination with genetic tdTomato SGN cytosol fluorescence (Ai14-Neurog1-PSD), the rendered SGN surface was used as a reference for the PSD-HaloTag channel. Ribbon precursors were detected with the IMARIS Spots function (0.5 seed point diameter; 1.0 PSF correction, background subtraction, 15 seed point quality threshold, 45 region border growing; lineage tracing, 1.5 μm maximum distance, 0 maximum time gap, tracks minimum length 3 frames). Particle tracking for the precursors was conducted semi-automatically, using the lineage tracing algorithm. The fusion and fission of precursor particles was then checked manually, and based on fluorescence intensity center and independent mobility, individual particles were added to or removed from the automatically rendered tracks.

Live ribbon precursor volumes were detected using the surface rendering function (0.1 surface detail, local background subtraction, 0.28 μm largest sphere diameter, 0.3 split surface seed points, 3 quality filter, filter closest distance to Spots=Ribbons). For pharmacologically-treated conditions, local background subtraction was set to 25. Ribbon precursor spots as well as surfaces were classified as either in close proximity to the rendered PSD surface (≤ 500 nm), thus synaptically-engaged, or non-synaptic. In context of the IHC membrane, non-synaptic ribbons were subdivided into membrane-proximal precursors (≤ 500 nm from the rendered membrane surface) or free-floating, cytosolic precursors. Ribbon precursor classes were assigned per time frame, upon which all individual precursor data points over time were pooled per class. The classification of precursor tracks was assigned according to the predominant localization of precursors within the total trajectory: i) tracks containing synaptically-engaged ribbon precursors in minimally three consecutive imaging frames were classified as ‘synapse-interacting’ or synaptic tracks; the non-synaptic tracks could, in context of the IHC membrane, be divided in two subcategories: ii) membrane associated tracks consisted of predominantly membrane-proximal ribbon precursors, whereas iii) cytosolic tracks exclusively contained precursors that resided in the cytosol or localized in IHC membrane proximity for less than three consecutive frames. Ribbon precursor surfaces, spots and tracks were controlled for atypical characteristics and intracellular localization, such as large protein aggregates or potentially disrupted tissue, and excluded from the analysis.

Track segments for the synaptic context to ribbon mobility correlation analysis were selected manually from the semi-automatically rendered traces, after removal of fusion and fission events. Here, segment selection was based on ribbon precursor location in respect to the rendered PSD surface, bordered by the fluorescent SGN: (i) segments classified as AZ-approaching contained precursors initially distanced from the PSD and a final three consecutive imaging frames localizing in engagement with the AZ; (ii) continuously synaptic segments excluded any time precursors spent outside of synaptic engagement of over three consecutive imaging frames; (iii) segments of ribbon precursors distancing from the AZ were required to contain initial localization in an synaptically engaged state of three imaging frames, after which precursors localized at a further distance from the labeled PSD. Importantly, the precursor location in the analyzed track segments was controlled for proximity to other PSD-HaloTag fluorescent clusters in addition to target PSD, to exclude for potential confounding factors for tracks interacting with multiple PSDs.

The assessment of three-dimensional ribbon precursor displacement within the track segments was performed as in Voorn et al. 2024. The mean squared displacement (MSD) was calculated based on the extracted xyzt ribbon precursor coordinates, rendered in the IMARIS Spots function by center of mass. The traversed displacement was then averaged over the progressive imaging frames (τ).

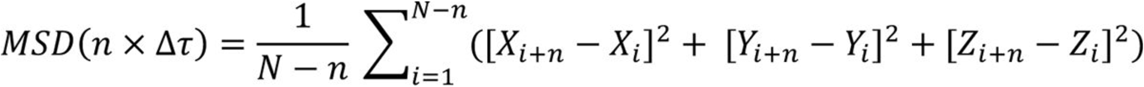

Here, x, y and z are the ribbon precursor coordinates, N is the number of data points in the track segment, and Δτ is the time interval per imaging frame. The MSD curve was then fitted using least squares fit to determine exponent α, as well as parameter K.

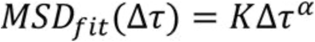

Track segment selection for ribbon precursors undergoing intersynaptic exchange was based on precursor location in respect to two different PSDs. Here, positional engagement was required with the primary AZ for three initial imaging frames, as well as thereafter with a secondary AZ for minimally three imaging frames. In cases of the two PSDs neighboring one another in such manner that the precursor trajectory remained in proximity to both PSDs while traversing, the length of the track segment was based on the relative distances between precursor and PSDs. From these segments, values such as displacement length, duration, mean and maximum velocity were extracted. For determination of precursors fusion at the secondary AZ, the subsequent continued track was also taken into account. The two PSDs involved in intersynaptic exchange were identified based on cohesive HaloTag fluorescence and respective mobility. Here, the volume, area and dynamics of singular PSDs within clearly defined fluorescent SGN boutons were exemplary in identifying a selection of smaller PSD puncta in close proximity as one postsynaptic contact. In instances where this assignment proved difficult, the total was regarded as one postsynaptic contact.

The intracellular location of ribbon precursors was classified as pillar or modiolar according to assigned coordinates by placement of an axis origin point at the IHC base in IMARIS (‘Reference Frame’). Based on the IHC orientation, pillar and modiolar sides could be distinguished. The axes of the Reference Frame were then positioned such that the x axis was parallel to the line of neighboring IHCs and the z axis was parallel to the positioning of the IHC *in situ*. For individual precursors, exact assigned coordinates were used for the analysis; for precursor tracks, the mean position over time was used.

### Statistics

Statistical analyses were performed in Prism8 (GraphPad, San Diego, CA). Values in the Figure descriptions are presented as classification_median_, classification_IQR_ (inter quartile range), or as mean ± SEM. For the pharmacologically-treated conditions, values were normalized against the distribution median of the matching classification prior to drug application. Normality of the distributions was determined using a D’Agostino-Pearson test. Statistical significance between two groups was determined with a Mann-Whitney U test; for comparison between multiple groups a Kruskal-Wallis test was performed in combination with a multiple comparisons Dunn’s post-hoc. Correlation between various parameters was determined using the Pearson’s R correlation matrix.

## Results

### Monitoring real-time ribbon dynamics *in situ*

To investigate the mobility and structural plasticity of IHC ribbon precursors, we established triple-color live-cell synapse labeling by crossbreeding various reporter mouse lines for cellular contexting of either the IHC plasma membrane – via YFP-tagged ChR2 expression in IHCs (Ai32-VC-KI-PSD); or the postsynaptic terminal – through cytosolic tdTomato expression in a small percentage of SGNs (Ai14-Neurog1-PSD) (Figure 1A-B). We then performed intra-cochlear AAV injections to introduce spectrally-matched RIBEYE-xFP and acutely applied fluorescent HaloTag ligands to visualize PSD95 (Fig.1C-E). Since continued viral overexpression may cause experimental artifacts, we opted for a short-term culture system, in which transduced IHCs were analyzed within 2-3 days post-injection. Here, ribbon counts and volumes were indistinguishable from wildtype ribbons ^14^, and RIBEYE-xFP colocalized with the presynaptic proteins Piccolino and Bassoon (Fig.1F-G). Next, we evaluated the labeling efficiency of different HaloTag ligands (JF646 and JF549) using post-hoc immuno-analysis with a PSD95-directed nanobody. Both ligands accurately labeled the PSD and displayed excellent colocalization with αPSD95 staining around the basolateral IHC compartment (Fig1.H), where Halo ligand fluorescence partnered with endogenous ribbons (Fig.1I).

**Figure 1.**
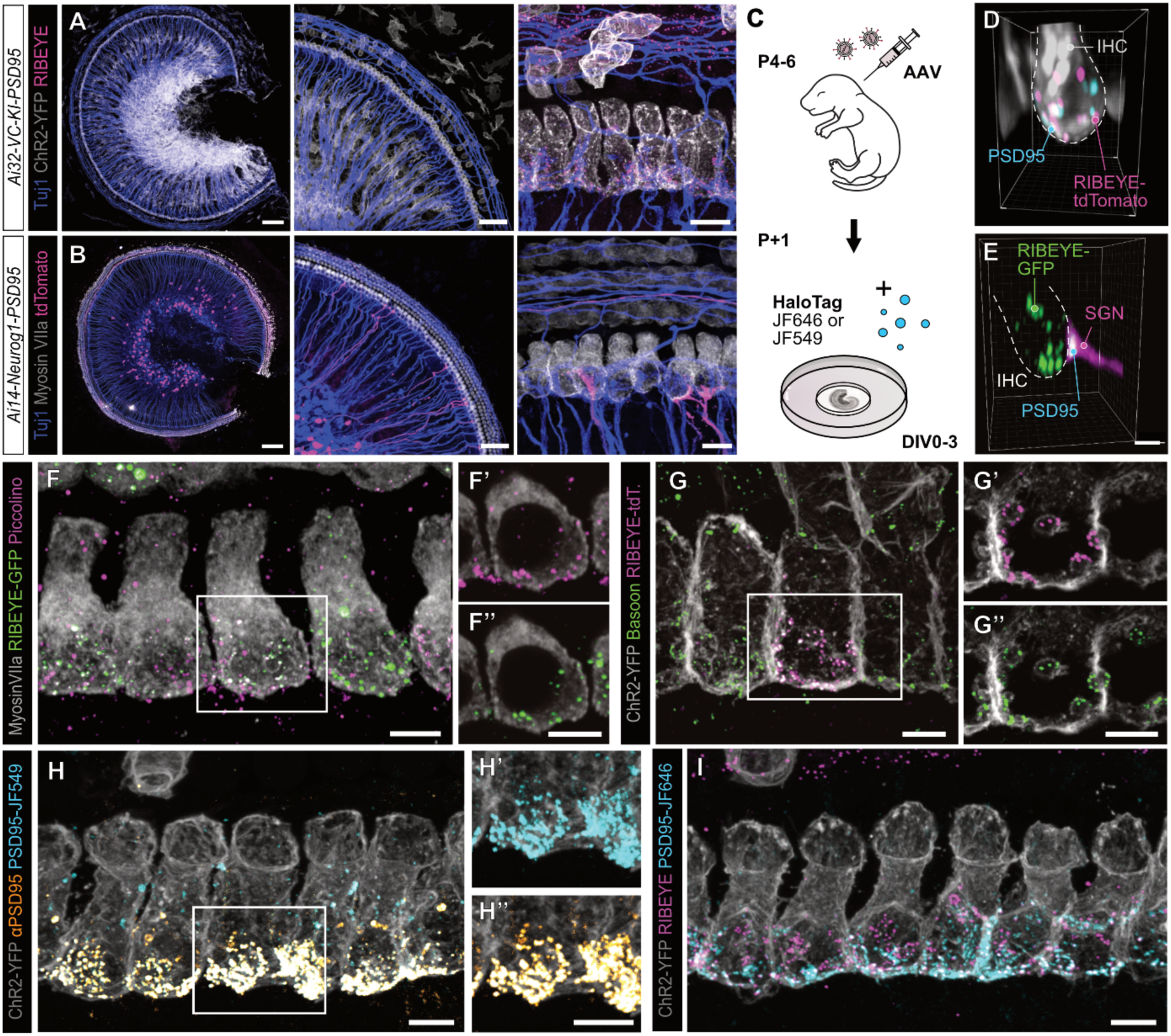
Experimental setup for the tracing of IHC ribbon precursor dynamics in neonatal mice. **A.** Tissue overview of an organotypic culture of the organ of Corti of an Ai32-VC-KI-PSD95 mouse with endogenous YFP fluorescence of the IHC plasma membrane. **B.** Tissue overview of an Ai14-Neurog1-PSD95 mouse with endogenous sparse labeling of SGNs with tdTomato. **C.** Experimental setup, making use of the mouse lines in A and B. Viral injections of a RIBEYE-XFP AAV (either -GFP or -tdTomato) are performed in neonatal mice, upon which the organ of Corti is extracted and organotypically cultured one day post injection. In vitro application of a HaloTag ligand (Janelia Fluor (JF) 549 or 646) to label HaloTag-linked PSD95 protein enables triple-color fluorescent labeling of cellular context, presynapse and postsynapse. **D, E.** The resulting live triple-color labeling described in C, for the mouse lines Ai3-VC-KI-PSD (D) and Ai14-Neurog1-PSD (E). **F, G.** Virally overexpressed RIBEYE colocalizes with presynaptic proteins Piccolino (F) and Bassoon (G); z-projections. **F’, F’’.** Isolated fluorescence of Piccolino (F’) and of RIBEYE-GFP (F’’), single plane of F. **G’, G’’.** Isolated fluorescence of RIBEYE-tdTomato (G’) and Bassoon (G’’), single plane of G. **H.** Application of PSD95-HaloTag ligands (H’) results in a labeling pattern highly comparable to PSD95 antibody staining (H’’), showing almost complete colocalization, especially in basolateral IHC compartments. **I.** The PSD95-HaloTag mediated labeling results in a near-native fluorescence pattern of postsynaptic PSD95 partnering presynaptic ribbon precursors. Scale bars: A, B left: 100 μm; middle: 50 μm; right: 10 μm; D-I: 5 μm.

### IHC ribbon synapse development is a dynamic process

Upon successful validation, we first studied IHC ribbon precursor dynamics in different intracellular fractions by longitudinally tracking tdTomato-labeled ribbon precursors within the basolateral IHC compartment. Due to the structural contexting, we could distinguish three distinct classes of ribbon precursors: (i) synaptically-engaged precursors, (ii) membrane-proximal, but non-synaptic precursors, and (iii) cytosolic precursors (Fig.2A) that could then be differentially analyzed for their respective mobility behavior (Fig.2B). Of the total precursor population, the majority registered as non-synaptic/membrane-proximal (∼60%), whereas ∼25% were synaptically-engaged, and another ∼15% cytosolically floating (Fig.2A). Interestingly, volume analysis revealed that the synaptic precursors were on average smaller than non-synaptic/membrane-proximal or cytosolic precursors (Fig.2C,D). Yet, they formed denser clusters than the membrane-proximal precursors, whereas cytosolic precursors were considerably more dispersed (Fig.2E).

**Figure 2.**
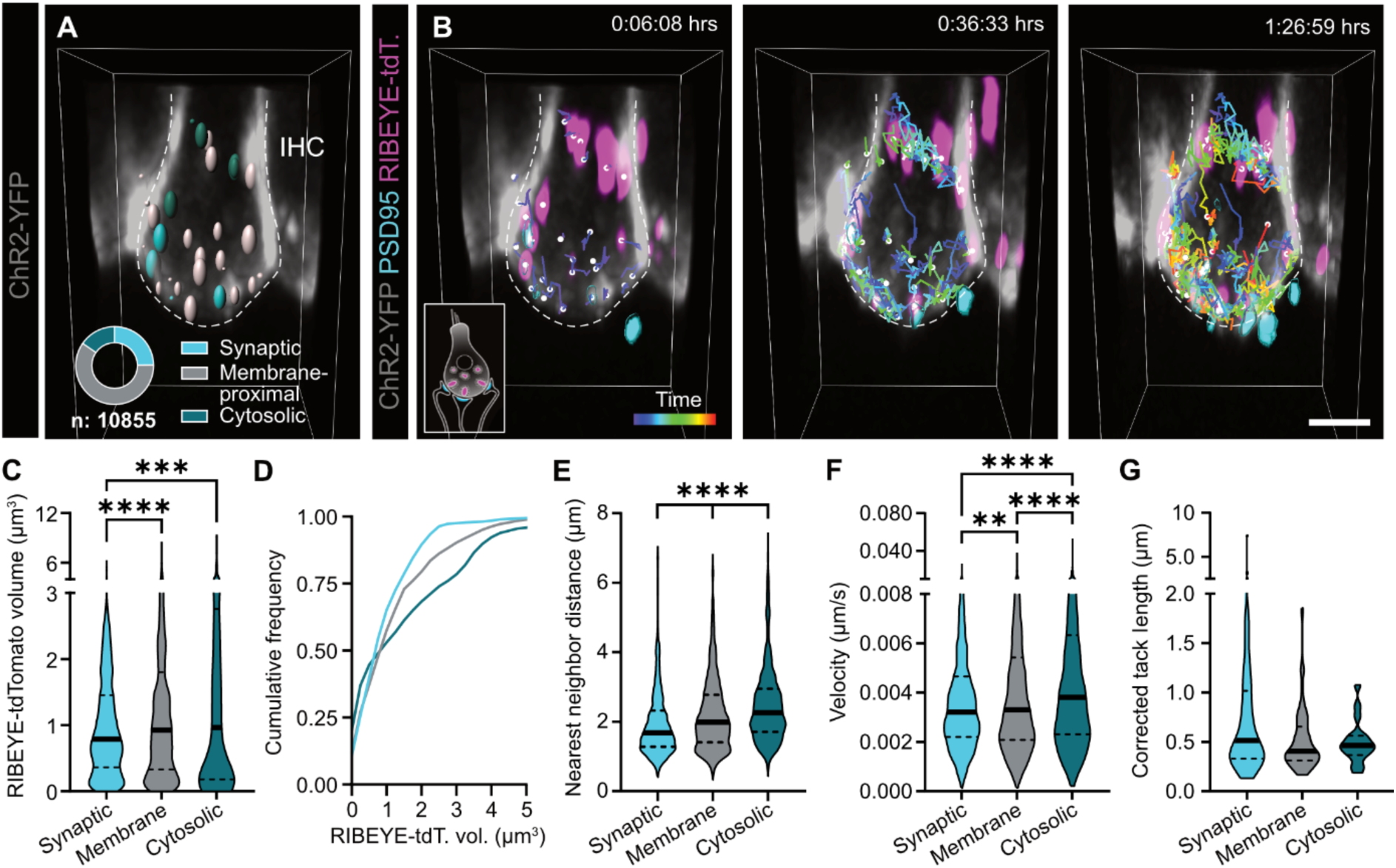
Ribbon precursor dynamics depend on the intracellular localization. **A.** Representative 3D image of an IHC expressing RIBEYE-tdTomato. Ribbon precursors are indicated with spheres, color coded for three different precursor classes based on their intracellular localization: synaptic, membrane-proximal, and cytosolic. **B.** Time-lapse imaging frames taken over 90 minutes, showing the progressive development and complexity of the ribbon precursor tracks, color coded for time. The insert illustrates the experimental setup. Frames taken from a triple-color live imaged organotypic culture of the organ of Corti (P5DIV2), with endogenous IHC membrane fluorescence, overexpression of RIBEYE-tdTomato, and the PSD labeled by PSD95-HaloTag ligand (JF646). **C.** The volume of synaptic ribbon precursors is lower than of membrane-proximal and cytosolic precursors. **D.** The frequency distribution of the different precursor classes shows a significant fraction of the cytosolic precursors consists of considerably larger precursors. **E.** The nearest neighbor distance (NND) of synaptic precursors is relatively low, while that of membrane-proximal precursors is slightly higher, and the NND of cytosolic precursors is much higher, indicating a more dispersed localization. **F.** The velocity of synaptic ribbon precursors is slightly lower than of membrane-proximal precursors, whereas cytosolic precursors appear to displace at a much higher pace. **G.** The length precursor tracks over the course of imaging time is not significantly different between the different classes, although the spread of the synaptic track length is considerably larger compared to that of the membrane-proximal and cytosolic precursors. **p<0.01, ***p<0.001, ****p<0.0001. Statistical significance: Kruskal-Wallis. N_animals_=4, n_IHC_=9, n_ribbons_=10855. Scale bar: 5 μm.

Next, we examined the dynamic characteristics of ribbon precursors across the acquisition time and found that cytosolic precursors displaced with the highest velocity, while membrane-proximal and synaptically-engaged precursors displayed increasing degrees of confinement (Fig.2F). This behavior is compatible with molecular trapping within the CAZ. Interestingly, this difference did not hold up when ribbon precursor velocity was averaged across the traveled trajectory: Here, despite the comparable track mobility, consideration of the full precursor track did show a remarkably larger spread in track length for synapse-interacting than non-synaptic or cytosolic tracks (Fig.2G), suggesting differential dynamics between AZ-associated precursors and the other fractions.

### Ribbon precursor plasticity centers around the developing AZ

Consistent with our previous work ^14^, long-term tracking of ribbon precursor dynamics yielded a complex network of interconnected trajectories that was characterized by numerous fusion and fission events (Fig.3A-B). Upon categorization, synaptic tracks appeared most complex and displayed the highest number of plasticity events of the different track classes (Fig.3C). However, surprisingly, fusion events were matched in frequency by fissions (i.e., plasticity events where a ribbon would *lose* material) thus rendering the bi-directional structural plasticity of ribbon precursors at nascent AZs strikingly balanced (Fig.3D). Moreover, when calculating the frequency distributions, synaptic tracks were most plastic and underwent several consecutive events of fission and re-fusion in our imaging window – apart from a small non-interactive subpopulation that solely underwent internal remodeling while continuously residing at the AZ (Fig.3E-F). While fusions clearly present a physiological mechanism of rapidly boosting ribbon volume (Fig.3G-G’), fissions likely serve to cap ribbon size and redistribute the material of an initially large ribbon precursor over multiple smaller secondary precursors – potentially with a secondary function in ‘seeding’ additional AZ scaffolds at novel locations (Fig.3H,H’). Consistent with previous work from zebrafish lateral line neuromast hair cells (nHC ^32^), we observed instantaneous appearance of diffraction-limited precursors in our observation volume indicative of *de novo* precursor assembly from a soluble RIBEYE pool. In further support of such a ‘cytoplasmic reservoir’ hypothesis, we detected a gradual volume increase in synaptic ribbons lacking detectable plasticity events (Fig.3I,I’).

**Figure 3.**
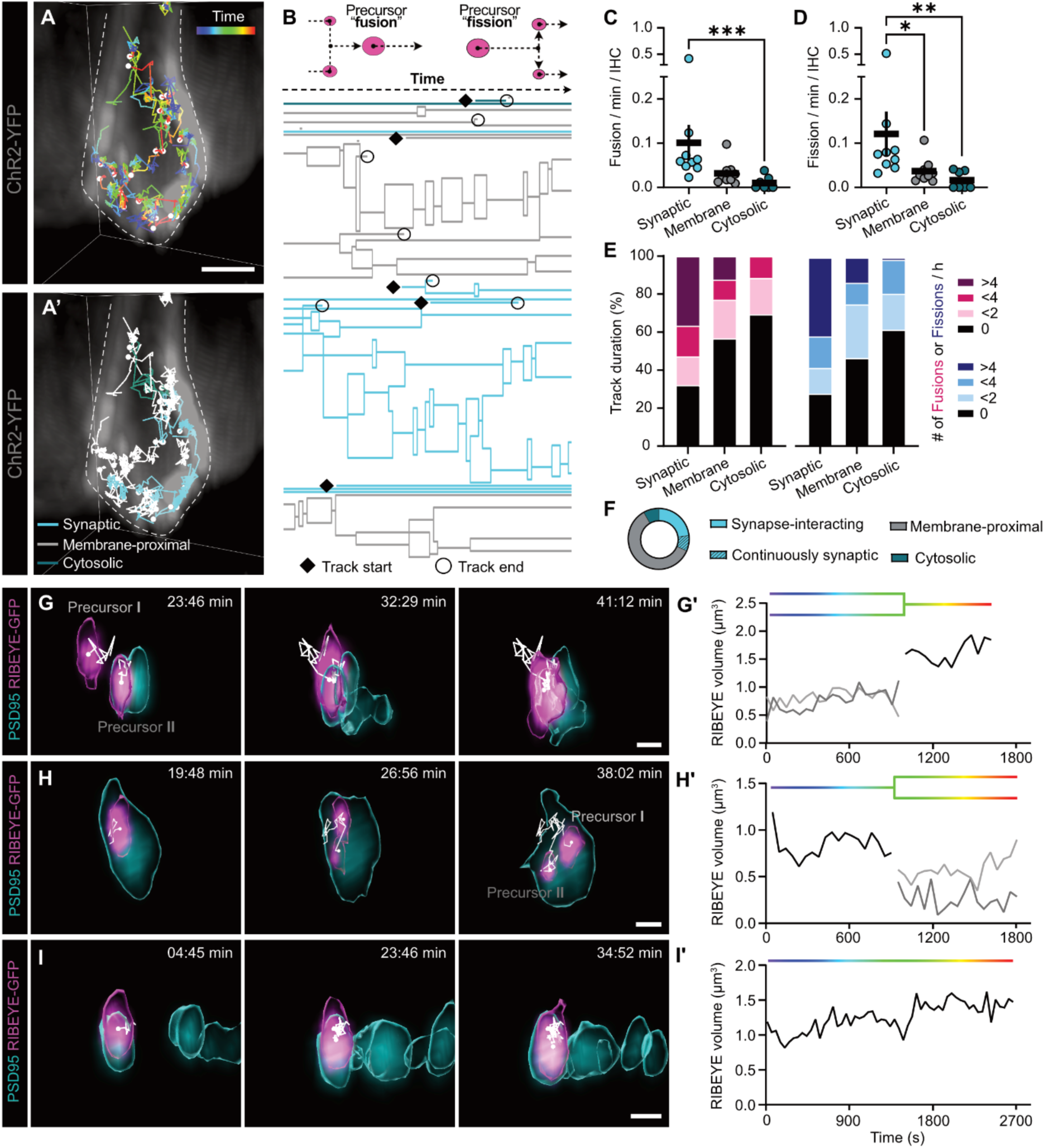
Ribbon precursor plasticity centers around the presynaptic AZ. **A.** Exemplary 3D live imaged frame of an IHC expressing RIBEYE-tdTomato (P6DIV2), with indicated ribbon precursor tracks over 90 minutes, color coded for time (A), and for the three different precursor track classes (A’): synapse-interacting, membrane-proximal and cytosolic. **B.** Lineage tracing graph illustrating ribbon precursor plasticity over time. Each line represents a single precursor; mergers and splits display fusion and fission events. Certain precursor tracks appear to start during imaging time (diamonds), whereas other tracks become undetectable (circles). **C, D.** Quantification of the plasticity event frequency shows that synapse-interacting precursor tracks contain more fusion events (C) than cytosolic tracks, and more fission events (D) than membrane-proximal, as well as cytosolic precursor tracks. **E.** The frequency distributions of precursor plasticity demonstrate the synapse-interacting tracks to be highly plastic. **F.** Distribution of the three different precursor track classes, according to the total sum of track duration. A small subpopulation of the synapse-interacting tracks consists exclusively of ribbon precursors that continuously localize at the presynaptic AZ. **G-I.** Exemplary 3D rendered time-lapse imaging frames, showing volume adaptation of ribbon precursors at the AZ during fusion (G), fission (H), or stable, non-plastic localization (I). **G’-I’** Volume progression over time of the precursors in G-I, color coding as indicated in G-I. Lineage tracing of precursor plasticity is indicated at the top of the graph, color coded for time. *p<0.05, **p<0.01, ***p<0.001. Statistical significance: Kruskal-Wallis. N_animals_=4, n_IHC_=9, n_ribbons_=10855. Scale bars: A: 5μm; G, H, I: 1 μm.

### Synapse location influences ribbon precursor plasticity

Several studies have detailed the functional heterogeneity of IHC synaptic contacts in mature IHCs. Here, location-dependent differences in synapse volume, voltage-dependence of Ca^2+^ influx, postsynaptic SGN caliber, excitability and RNA profiles give rise to a pillar/modiolar gradient of low to high threshold synapses ^16,17,33,34,53^. While the postsynaptic diversification has been suggested to originate in the earliest developmental stages and requires pre-sensory SA, the mechanisms underlying presynaptic heterogeneity and its temporal sequence remains unclear. However, recent findings demonstrate early differentiation between low vs. high spontaneous rate (SR) SGN profiles along the pillar/modiolar IHC perimeter of young postnatal mice *in vivo* ^34^. We thus probed if location-dependent differences in ribbon dynamics could also be detected in our live-cell imaging paradigm and mapped individual ribbon precursor trajectories onto the pillar/modiolar-cuticular/habenular IHC axes (Fig.4A,B). We then performed location-based comparative analyses of all precursors, separated by a hard midline “0” cutoff between the pillar/modiolar IHC halves to observe global effects. To refine this analysis, we then excluded the subpopulation with intermediate coordinates and selected for the precursors at the pillar and modiolar ‘extremes’, which are poised to give a clearer picture on low-threshold, high SR vs. high-threshold, low SR fibers (Fig.4B’). Here, the global trends observed in the basic midline pillar/modiolar distinction were strikingly exacerbated in the selective populations, with the > ±3 μm cutoff offering the clearest representation of these effects without compromising the statistical power of our analysis (for additional cutoffs, please refer to Supp. Fig.S1). While individual ribbon volumes were lower on the modiolar side (Fig.4C), these ribbons clustered at lower nearest-neighbor distances (NND) (Fig.4D). We then investigated the dynamic properties of ribbon precursors and, while we found no location-dependent variations in velocity (Fig.4E), the level of ribbon precursor plasticity significantly differed between the IHC extremes, as reflected by a higher frequency of fusion and fission events in the modiolar tracks (Fig.4F-H). Upon closer inspection of the precursor subpopulation characteristics, we additionally found that modiolar ribbon volumes were lower across all precursor classes and NND only differed between the cytosolic precursor fractions (Fig.4I-J). For completion, we also assessed precursor dynamics and found no difference in precursor velocity (Fig.4K), but a notable trend towards lower modiolar fusion frequency in synaptic tracks (Fig.4L).

**Figure 4.**
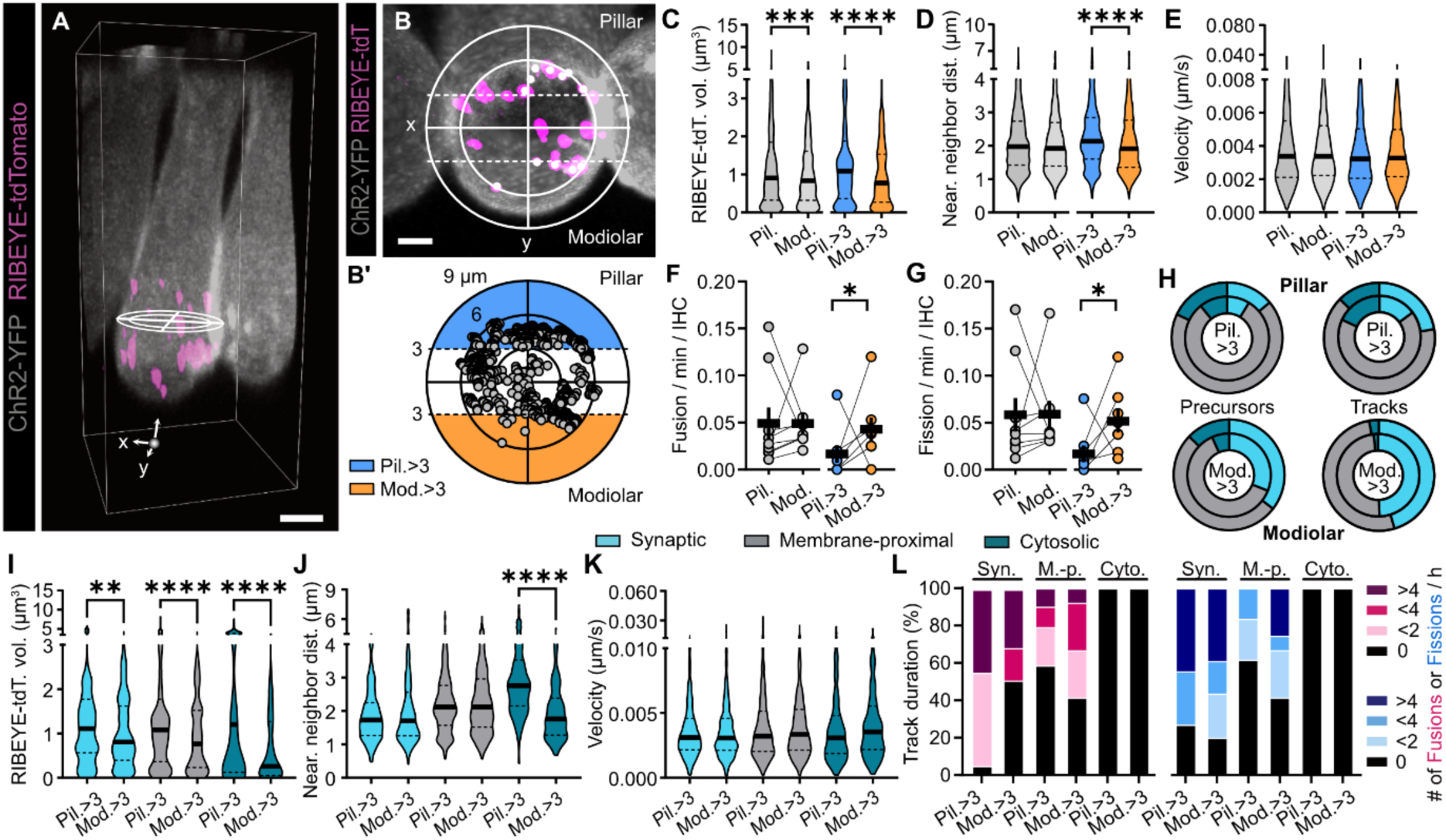
Ribbon precursor plasticity differs along the pillar/modiolar IHC axis. **A.** Exemplary 3D live image of an IHC expressing RIBEYE-tdTomato, with x and y axes indicated used to determine pillar and modiolar IHC fractions. **B.** Top-down view of A. Dotted lines demonstrate the pillar/modiolar extremities of the IHC >3 μm from the 0 midline. **B’.** Illustration of the >3 μm extremities, displaying all precursors over time. **C-G.** Pillar/modiolar populations based on the 0 midline (grays), and the >3 μm division (blue/orange). **C.** The volume of modiolar precursors is lower than of pillar precursors. **D.** In the >3 μm extremities, the nearest neighbor distance (NND) is lower for modiolar than pillar precursors. **E.** Ribbon precursor velocity is comparable throughout the IHC. **F, G.** The frequency of fusion (F) and fission (G) events is increased in modiolar tracks in the >3 μm extremities. **H.** The fraction of synaptically-engaged precursors and tracks is considerably higher in the modiolar than the pillar IHC fraction, for the total (outer ring) and >3 μm populations (inner ring). **I-K.** Displaying solely the >3 μm populations. **I.** The volume of modiolar precursors was lower than of pillar precursors in all precursor classes. **J.** The lower modiolar NND appears to be driven by the cytosolic precursors. **K.** Precursor velocity was similar for all classes. **L.** The plasticity distribution shows a slightly higher frequency of fission events in modiolar synaptic tracks. *p<0.05, **p<0.01, ***p<0.001, ****p<0.0001. Statistical significance: Kruskal-Wallis. N_animals_=4, n_IHC_=9, n_ribbons_=10855. Scale bars: A: 5 μm; B: 2 μm.

Overall, these data suggest that location-dependent differences can already be observed during early postnatal development and may pose the developmental foundation for planar synapse heterogeneity in mature IHCs.

### Synaptic engagement restricts ribbon precursor mobility

We next shifted our focus from the entire basolateral IHC compartment to individual synaptic contacts and employed the *Ai14-Neurog1-PSD* line, in which sparse labeling of individual SGNs allowed us to trace ribbon dynamics specifically at the PSD of a singular afferent bouton (Fig.5A). We first traced all RIBEYE-GFP labeled precursors proximal to the fluorescent SGN, and then selected for AZ-interacting tracks. Intriguingly, individual ribbon precursors displayed various distinct mobility patterns along their trajectories (Fig.5B): for example, while the AZ approach often occurred with large displacement and high velocity, both parameters drastically decreased upon PSD engagement and continued residence at the AZ (Fig.5B’). Here, the apparent ‘capture’ of the precursor in the CAZ network did however not completely halt its mobility, but confined its movements (Fig.5C,C’). Notably, we made multiple observations of AZ-engaged precursor mobility being mirrored by the associated PSD (Supp. Fig.S2) – indicative of trans-synaptic coupling mechanism during synaptogenesis.

**Figure 5.**
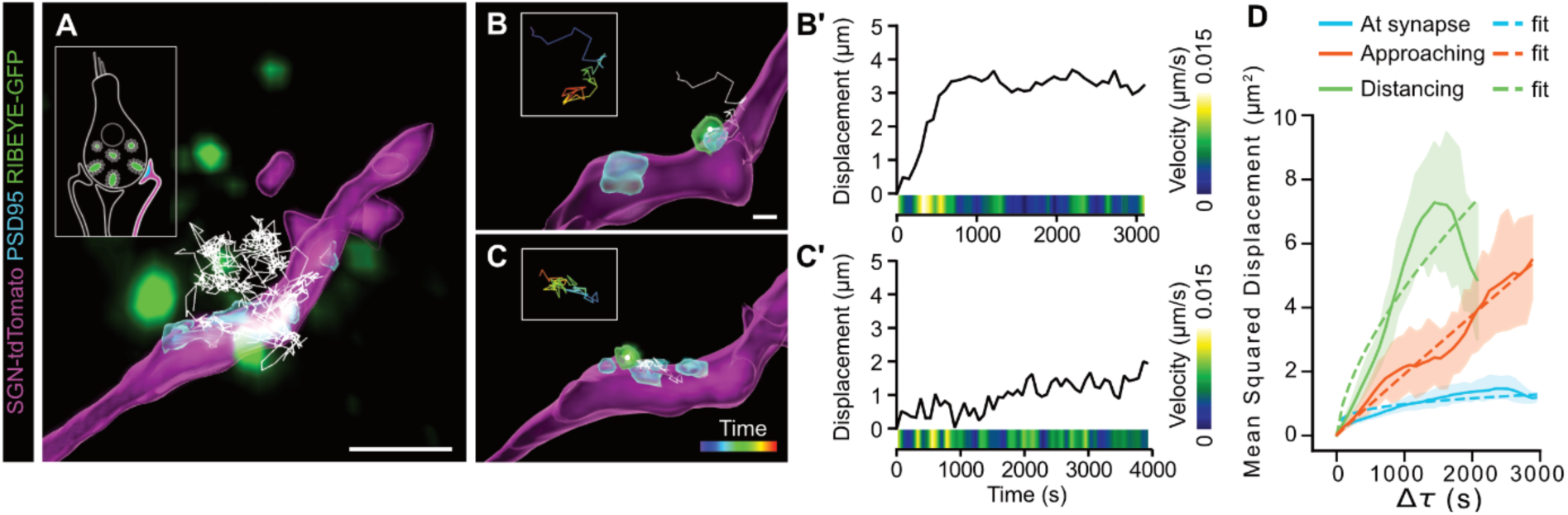
The context of the presynaptic AZ confines ribbon precursor mobility. **A.** Exemplary 3D reconstruction of a tdTomato-positive SGN with the PSD labeled by PSD95-HaloTag ligand (JF646), rendered from a live triple-color imaged organotypic culture of the organ of Corti (P6DIV1). The SGN innervates a RIBEYE-GFP expressing IHC, of which the traced ribbon precursor trajectories are indicated that interact with the synaptic contact of the SGN over 90 minutes of time-lapse imaging. The insert illustrates the experimental setup. **B.** Trajectory segment of a ribbon precursor that has displaced/traversed in approaching motion towards the SGN, and now localizes apposed a PSD inside the SGN. **C.** Trajectory segment of a ribbon precursor continuously localized in engagement with the PSD inside the SGN, therefore likely continuously at the AZ. **B, C.** Inserts: trajectory segment color coded for time. **B’, C’**. The displacement length and velocity (color coded) of the ribbon precursors in examples B and C over time, showing a steep rise in displacement and peak in velocity when (B’) the ribbon precursor approaches the synapse. **D.** The mean squared displacement (MSD) of three different trajectory segment classes: precursors in motion of AZ approach (orange), distancing from the AZ (green) and in continuous association with the AZ (cyan). The displacement of precursors at the AZ is confined compared to more mobile precursors in distancing or approaching motion. At synapse, α=0.24; Distancing, α=0.64; Approaching, α=0.97. N_animals_=7, n_IHCs_=12, n_segments_=18. Scale bars: A, 5 μm; B, C: 1 μm.

To now differentially quantify the interplay between ribbon precursor mobility and AZ engagement, we further analyzed stable track segments and, based on their position, categorized them as: (i) AZ-approaching, (ii) continuously synaptic, or (iii) AZ-distancing. We then plotted the mean squared displacement (MSD) for the different track segments, and fitted the average per interaction type (Fig.5D). We found that – compatible with a mechanism of diffusional trapping – displacement of the continuously synaptic precursors was strongly confined, whereas the mobility of the AZ-approaching and AZ-distancing precursors contained distinctly higher displacements, thus implicating the participation of likely different molecular motor-based transport mechanisms.

### Ribbon precursors undergo intersynaptic exchange between AZs

Since the sparse SGN labeling approach allowed us to delineate between PSD fragments within a given terminal and neighboring PSDs of nearby fibers (Fig. 6A), we could analyze the frequency of intersynaptic exchange events. Here, we observed precursors detach from established synaptic contacts, or fission events from synaptically-engaged ribbons that resulted in transfer of ribbon material to a secondary PSD. Although challenging to quantify, we estimated intersynaptic exchange to occur at ∼25% of all traced postsynaptic sites (Fig.6A-B). However, this might be a cautious estimate due to our conservative assignment of individual postsynaptic contacts in cases of more ambiguous identification without fluorescent SGN context label. Of the sites displaying intersynaptic exchange, the frequency of occurrence was generally low (Fig.6C), with mostly single plasticity events presenting during our observation window. In few cases, multiple events could be detected, where a single site could be both donating and accepting ribbon material, or have precursors detach and re-attach in the same location. In ∼50% of cases, intersynaptic exchange events were followed by fusion of the exchanged material with a membrane-resident synaptic ribbon at the secondary postsynapse (Fig.6B’,E-F, Supp. Video 1+2). While the traveled distance and duration of exchange varied considerably between individual events, it displayed a positive – yet statistically non-significant – trend (Fig.6D). Notably, most of these events occurred between directly neighboring synapses and where hence short distanced; however, several instances covered multiple micrometers (range: 0.56 – 7.35 μm), with ribbon precursors translocating across the entire basolateral IHC pole in a surprisingly targeted manner that followed a near-linear trajectory (Fig.6E). Interestingly, we also observed instances of seemingly targeted slow precursor ‘walking’ motion or mixed modes (Fig.6F-G’). Regardless of the translocation mode, closer inspection of the velocity profiles of detaching precursors revealed an initial velocity peak upon detachment from the primary ribbon/AZ, indicative of an energy barrier that needed to be overcome for successful fission.

**Figure 6.**
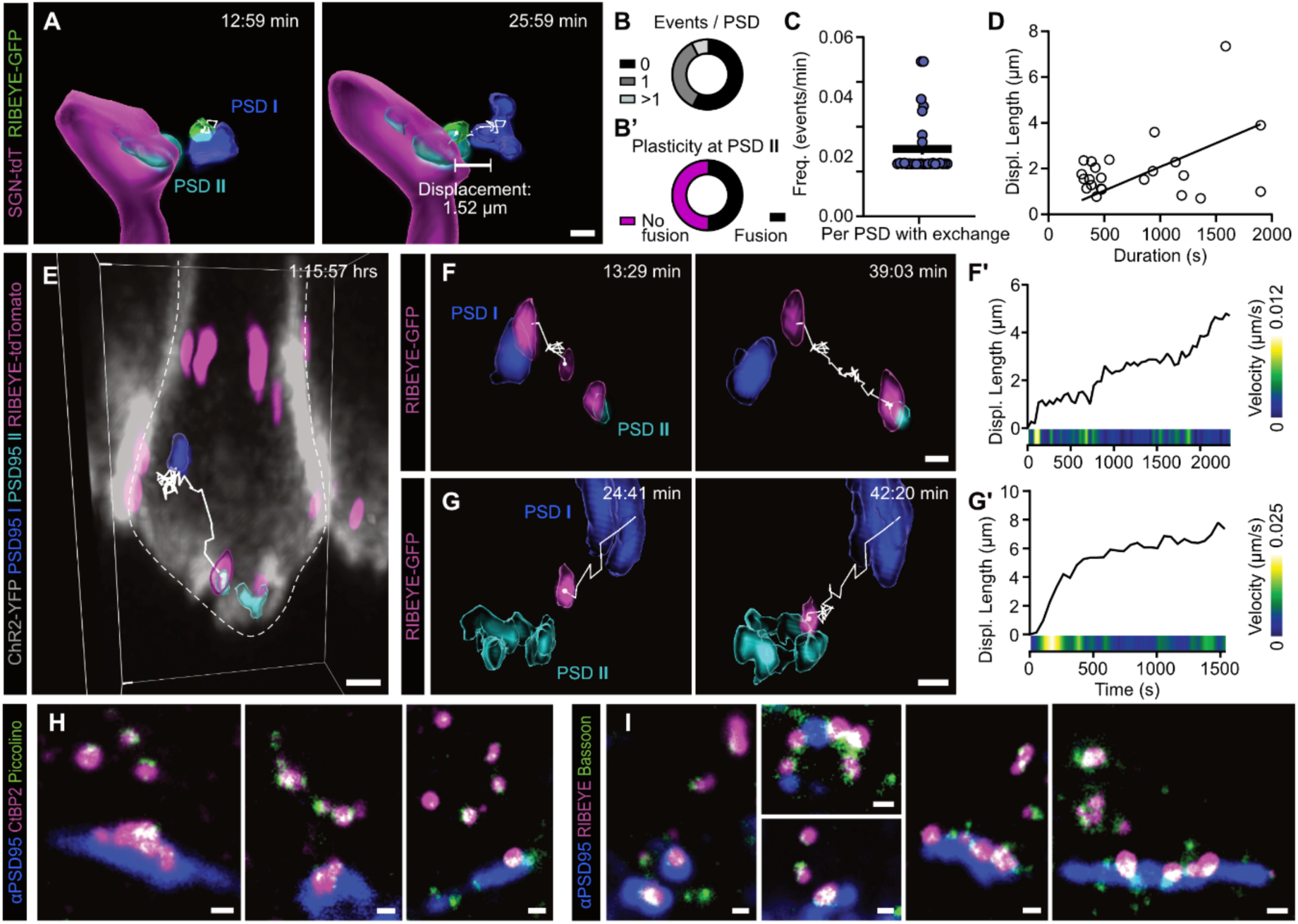
Intersynaptic exchange of ribbon precursors takes place between different presynaptic AZs. **A.** Illustrative example of intersynaptic exchange between a primary synaptic contact (dark blue PSD) situated outside and a secondary synapse (cyan) inside the fluorescent SGN; rendered from a triple-color live imaged organotypic culture of the organ of Corti (P7DIV1) **B.** Frequency of events of intersynaptic exchange per traced postsynapse. **B’**. Frequency of ribbon precursor fusion at the secondary synapse. **C.** Frequency of events per PSD for which intersynaptic exchange was detected. **D.** The displacement length of intersynaptic exchange trajectories showed a positive, but non-significant correlation with the trajectory duration. Y = 0.002198*X, p= 0.6769. **E.** Example of traced intersynaptic exchange of a ribbon precursor along the IHC membrane. **F.** Example of intersynaptic exchange where a small precursor can be observed to bud off from the larger precursor engaged at the primary synapse. During the exchange, the synaptically-engaged precursor uncouples from the primary PSD. **G.** Example of intersynaptic exchange of a precursor traversing a long distance between the two PSDs, with an initial rapid distancing from the primary synaptic contact. **F’, G’.** Displacement length and precursor velocity (color coded) over time of examples in F and G. **H, I**. STED microscopic images of synaptically-engaged and non-synaptic precursors, showing colocalization of (H) RIBEYE (labeled by CtBP2) with Piccolino in precursors in both intracellular locations, as well as (I) colocalization between RIBEYE and Bassoon. N_animals_=10, n_IHC_=14, n_traces_=23. Scale bars: A, E, F, G: 2 μm; H, I: 500 nm.

Since ribbon size has a direct impact on the SV tethering and CaV clustering capacity of a given AZ, we next employed STED nanoscopy to interrogate the molecular identity of AZ-proximal, cytoplasmically floating ribbon precursors (Fig.6H,I). Consistent with previous reports ^13,35^, we identified the ribbon-specific protein Piccolino as an essential structural component, albeit at vastly varying quantities. Moreover, the ribbon-anchoring protein Bassoon consistently co-localized with synaptically-attached, but also free-floating precursors. This suggests that the dynamic plasticity of the maturing AZs involves the transfer of functional ribbon building blocks, which could rapidly – and reversibly – modify exocytic output of a given AZ.

### Pre-sensory spontaneous activity drives ribbon plasticity

Lastly, we investigated the influence of pre-sensory SA on developmental ribbon dynamics. Considering the plastic changes in ribbon precursor size that have been described in young nHCs upon manipulation of presynaptic Ca^2+^ influx ^36^, we suspected that pre-sensory SA may also play a regulatory role in mammalian ribbon development. Therefore, we utilized the intrinsic SA retained in organotypically-cultured organs of Corti ^15,37^ and first assessed if acute SV release bouts would trigger ribbon displacement due to excessive AZ membrane turnover. Here, we correlated ribbon dynamics with postsynaptic activation using GCaMP6-triggered averaging of velocity traces, but failed to detect reproducible patterns – thus suggesting statistical independence of these parameters (Supp. Fig. S3). We then pharmacologically blocked presynaptic Ca^2+^ influx with the L-type Ca_v_ channel antagonist isradipine – thereby effectively silencing IHC signal transmission – and performed sequential time-lapse imaging before and after isradipine or vehicle (Fig.7A-C). Remarkably, isradipine strikingly reduced trajectorial complexity of synaptic tracks: When quantifying the frequency of plasticity events, the occurrence of both fusions and fissions, was significantly reduced upon isradipine exposure, whereas the vehicle group remained unaffected. Similarly, non-synaptic tracks displayed unaltered behavior, regardless of treatment. When plotting the frequency distributions of individual plasticity events (Fig.7D), fusions and fissions within the synaptic tracks were starkly reduced to a level comparable to non-synaptic tracks, hence indicating that presynaptic Ca^2+^ entry is indeed a strong driver of structural adaptation at IHC AZs.

**Figure 7.**
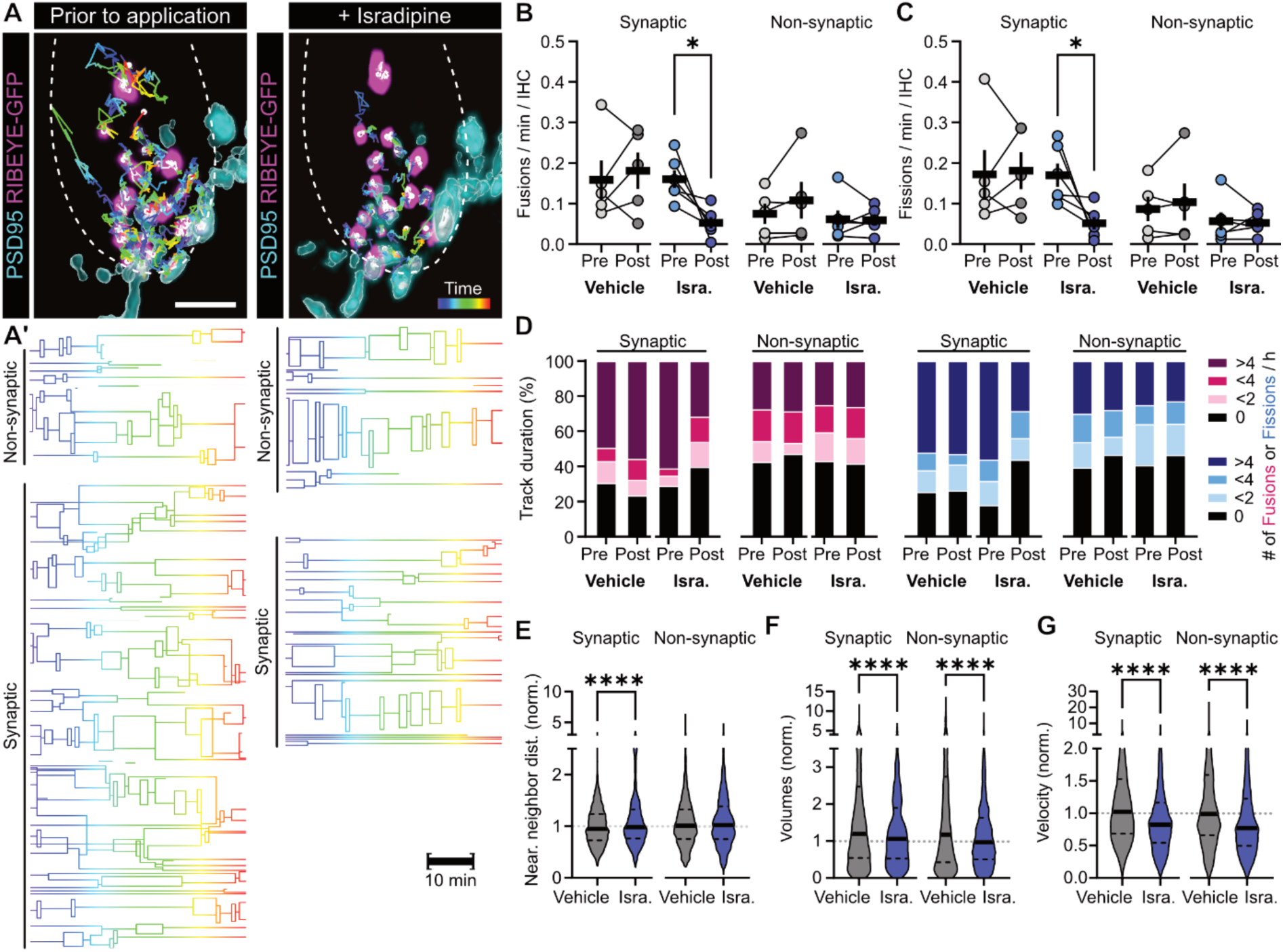
Inhibition of presynaptic Ca^2+^ influx reduces ribbon precursors plasticity around the AZ. **A.** Exemplary ribbon precursor traces over 45 minutes of an IHC expressing RIBEYE-GFP prior and post Isradipine application; PSDs are labeled by PSD95-HaloTag ligand (JF549), rendered in 3D from a live dual-color imaged organotypic culture of the organ of Corti (P7DIV1). **A’.** Lineage tracing graphs prior and post Isradipine application show the dramatic reduction in complexity of the synaptic precursor traces in A. **B, C.** The frequency of fusions (B) and fissions (C) upon Isradipine treatment is reduced in synaptic precursor tracks, whereas non-synaptic tracks are unaffected. Values displayed are pre and post pharmacological application, compared to a vehicle control (DMSO). **D.** The frequency distribution of plasticity events shows that the profile of synaptic tracks is reduced to the level of non-synaptic tracks upon exposure to Isradipine, for both fusion events (magenta) and fission events (blue). **E.** The nearest neighbor distance of ribbon precursors at the synapse is unaffected by Isradipine, but is slightly decreased in control conditions. **F.** A small increase in precursor volume is present in the control condition, while this is less prominent in synaptically engaged precursors, and absent in non-synaptic precursors in the Isradipine treated condition. **G.** Isradipine application reduced ribbon precursor velocity, whereas the control condition remained unchanged. **D, E, F.** Values displayed are post pharmacological treatment, normalized against the median of the paired precursor class prior to treatment. ****p<0.0001. Statistical significance: Kruskal-Wallis. N_animals_=7, n_IHC_=11, n_ribbons_=42456. Scale bar: 5 μm.

To extend this analysis, we examined the effects of isradipine on individual precursor level. We detected small changes in the NND and volume in untreated IHCs, which indicate a progression towards larger ribbons, and denser clustering at the synapse over time. Here, isradipine appeared to hinder this progression, resulting in unchanged NND (Fig.7E) but limited volume increase (Fig.7F). Finally, isradipine treatment significantly reduced precursor velocity – for both, synaptic and non-synaptic precursors (Fig.7G) – whereas the velocity in control conditions remained unchanged. Together, these data suggest that the loss of presynaptic Ca^2+^ influx attenuates ribbon precursor mobility, maturation and plasticity around the synaptic AZ, and may thereby play a prominent role in ribbon synapse development.

## Discussion

In the present study, we developed an *in situ* multi-color live-cell imaging approach to monitor mammalian IHC ribbon synapse maturation. We found that during development, ribbon precursors display a high degree of location-dependent spatial mobility and bi-directional structural plasticity (Fig.8). Ribbon precursors translocate in complex interactive networks centered around the presynapse, which captures, confines and exchanges precursors between adjacent AZs – thereby enabling rapid AZ formation at novel sites or structural supplementation of established synapses. We further explored the effects of pre-sensory SA on ribbon dynamics and identified activity-dependent Ca^2+^ influx as a key driver of structural plasticity at auditory ribbon synapses.

**Figure 8.**
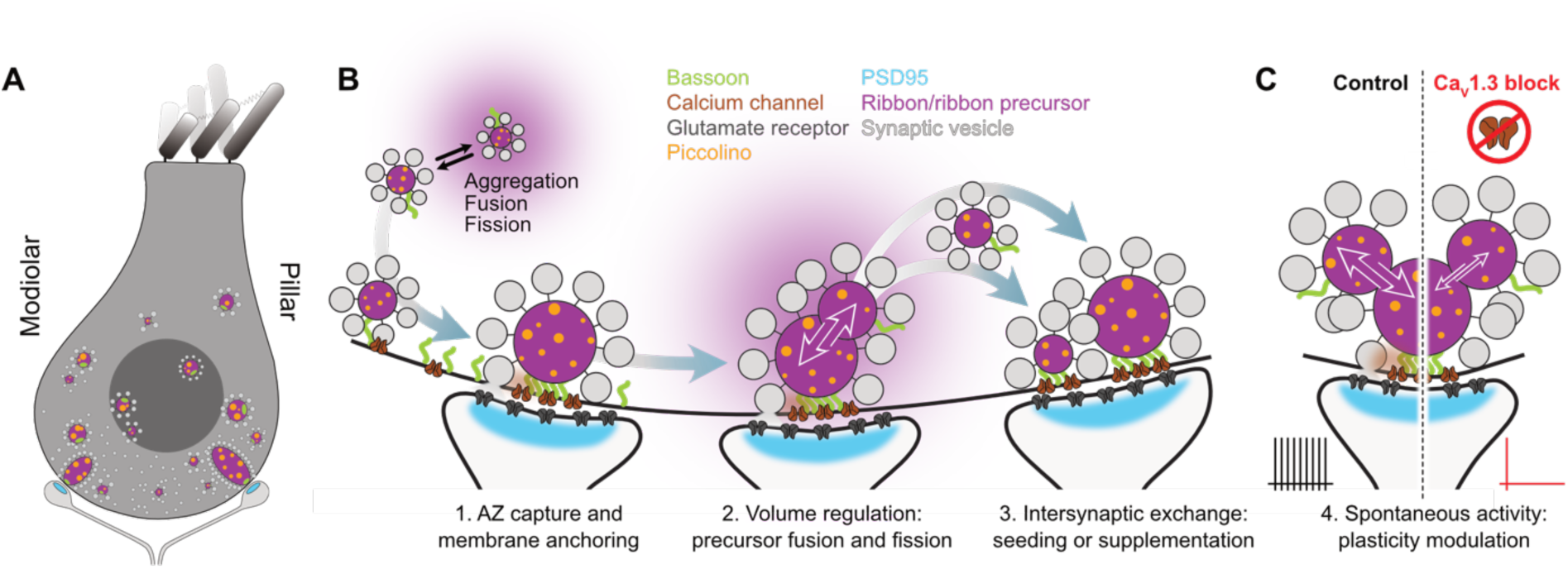
Overview of IHC ribbon synapse assembly in early postnatal development. **A.** The structural plasticity of ribbon precursors differs along the pillar/modiolar IHC axis: The modiolar ribbon fraction contains precursors that are of lower volume and more plastic than pillar precursors, and cluster at a closer distance in the cytosol. Also, the modiolar population contains a higher percentage of synaptic precursors and tracks. However, there is no pillar/modiolar difference in precursor number or velocity. **B.** Ribbon synapse assembly in IHCs involves three main processes: 1. The engagement of ribbon precursors with the active zone (AZ), through the immobilization and capture of precursors at the AZ, as well as the anchoring to the AZ plasma membrane; 2. The regulation of ribbon precursor volume through events of precursor fusion and fission, in the cytosol (1), as well as at the AZ (2) and additional condensation of soluble RIBEYE from a cytoplasmic pool at the ribbon scaffold and/or fusion of small diffraction-limited precursors. Ribbon precursor aggregates of all detected volumes consisted of RIBEYE, Piccolino and Bassoon – hence implying the transfer of complete organelle building blocks; 3. The exchange of ribbon precursor material between different presynaptic AZs, to either reduce or supplement existing synapse, or ‘seed’ new AZs. **C.** Presynaptic Ca^2+^ influx appears to be a crucial factor in the developmental plasticity of IHC ribbon precursors, since the blocking of presynaptic Ca_V_s significantly reduces ribbon precursor plasticity at and in proximity of the presynaptic AZ.

### Developing ribbon-type AZs are structural plasticity hubs

Based on previous work using ‘static’ imaging methods ^11–13,38^ auditory ribbon synapse assembly has been suspected to be a uni-directional process, in which small ribbon precursor spheres self-assemble in the cytosol, gradually accumulate near the AZ to aggregate into larger complexes and eventually form mature, membrane-anchored ribbons ^11,13,18,38,39^. The present live-cell study challenges this simplistic view by revealing the balanced occurrence of precursor fusion and fission events that regulate individual ribbon size in a fluid manner. Importantly, structural plasticity required an intact synaptic context for maximum efficiency. Here, the AZ appears to fulfill two main functions: (i) precursor recruitment and (ii) stabilization for volume regulation and functional maturation. Upon engagement with the AZ, cytosolic precursors are spatially confined, adequately spaced and structurally adapted. Small precursors are regularly exchanged among membrane-resident ribbons and/or precursors that are not (yet) synaptically-engaged, thus leading to dynamic fluctuations in precursor volume, and contributing to the gradual refinement of ribbon shape towards hearing onset ^11–13^. In concert with such bulk volume adaptations, our findings indicate diffusional addition of soluble RIBEYE to already existing scaffolds, since structurally stable ribbons gradually acquire volume over time. To date, it remains unclear whether this represents separate regulatory pathways or rather reflects fusions and fissions taking place on a scale beyond the resolution limit of our acquisition system. Future use of photo-convertible fluorophores my help clarifying this issue.

### Ribbon precursor recruitment to the AZ

During AZ assembly, ribbon precursors undergo various types of mobility that – on first sight – appeared to follow a diffusion-like pattern reminiscent of Brownian motion. However, during their approach towards the AZ, ribbon precursor translocation was characterized by displacement and velocity peaks suggestive of targeted motion. This implies the involvement of rapid active transport via molecular motors, as reported for AZ-directed transport at conventional synapses ^40^. However, in contrast to axonal synapse assembly, the stretches of presynaptic protein displacement detectable in IHCs are considerably shorter and of multidirectional nature, which complicates the identification of cytoskeletal transport routes. Yet, the variety of ribbon precursor motion types aligns well with the multi-directional MT meshwork of IHCs ^14^. Moreover, considering the proximity of MT bundle endings to membrane-attached ribbons, their presumed +end polarity towards the AZ, colocalization of Kif1a to cytosolic ribbons and reduced ribbon volume accumulation upon Kif1a functional impairment ^13,14,41^, the IHC MT network appears to provide key transport routes for ribbon precursors translocation towards the AZ. In fact, MT-based trafficking facilitates precursor fusion and fission, as destabilization of the IHC MTs mainly affected the frequency of plasticity events ^14^. However, as pharmacological disruption of MTs did not entirely halt precursor displacement, additional contributions of other – e.g., actin-based – molecular transport pathways can be expected in this context and present interesting targets for future investigations.

### Ribbon precursor confinement at the AZ

Compatible with previous work on retinal photoreceptors ^42^, our data suggests that the CAZ serves to constrain precursor mobility. Here, the dense precursor clustering ensures sufficient spatial proximity for CAZ/ribbon interactions to take place and – at least transiently – limit ribbon precursor displacement. Such associations likely involve the common suspects Bassoon, Ca_V_1.3, RIM2, RIMBP2, and Piccolino (reviewed in ^43^). Interestingly, we found Piccolino and Bassoon to associate with all AZ-proximal precursors, regardless of their specific engagement status with the AZ. Hence, our findings suggest that IHC ribbon precursors serve as ‘AZ building blocks’ prior to AZ attachment, or during intersynaptic exchange. As an essential part of such building blocks, the presence of Bassoon suggests that floating precursors can attach at the plasma membrane easily and thus ‘seed’ ribbons at nascent AZs. However, considering the herein observed transient nature of ribbon precursor contacts with the IHC plasma membrane, sustainable anchoring may require additional post-translational modifications or other currently unknown CAZ interactions. Alternatively, as observed for conventional synapses, trans-synaptic pairing of cell adhesion molecules – including Neurexins and Neuroligins – may establish focal assembly points for adequate synapse formation ^44^. Indeed, our observations of paired ribbon/PSD95 cluster mobility indicate that trans-synaptic coupling between IHCs and SGNs plays a role in this process. Particularly, the Neurexin-Neuroligin connection has presented itself as an interesting candidate mechanism supporting ribbon anchoring and maturation in zebrafish nHCs ^45^ and mammalian IHCs ^46^. Adding to the complexity, various other cellular adhesion and ECM proteins have been localized to the IHC-SGN synapse ^47^, but remain to be investigated.

### Ribbon precursor dynamics differ along the pillar/modiolar IHC axis

The subcellular location of a given ribbon synapse along the pillar/modiolar IHC axis dictates its structural and functional properties ^48^. However, the underlying developmental mechanisms remain largely elusive. To address this, we examined location-dependent ribbon precursor dynamics in respect to this morphological axis. Interestingly, modiolar precursors displayed more ‘synaptic’ characteristics, with denser clustering, smaller individual volume, and higher frequency of plasticity events. Accordingly, a larger percentage of modiolar precursors and their associated tracks interacted with nascent AZs than in pillar locations. These findings are hence compatible with reports of modiolar synapses presenting with larger and more numerous synapses, as well as the retention of multi-ribbon AZs into adulthood ^13,33,49,50^. However, we detect a smaller individual volume of modiolar ribbons, whereas previous studies on developing IHCs found an inverse relationship between pillar and modiolar ribbons ^51,52^. Here, the tight clustering density we observed may have led earlier investigators to classify multiple ribbons as singular – thus larger – entities, especially as fixation-dependent tissue shrinkage may induce artificial clustering that would not have occurred in our live-cell analyses. Therefore, our dynamic data may provide a more realistic representation than chemically-fixed single time-point snapshots. Future experiments will be required to resolve this conundrum.

### Intersynaptic exchange facilitates AZ maturation

In the context of ribbon anchoring, the frequency of precursor dissociation from the AZ is surprisingly high and often coincided with a peak in precursor velocity. This suggests that a significant energy barrier needs to be overcome to detach precursors from its AZ-anchor or parent ribbon. Here, active processes are likely required to overcome such energy thresholds, as has been shown for the fission of other cellular organelles ^53,54^. Future studies will be required to identify the involved molecular cues and motors.

Such detaching precursors integrated into the highly plastic precursor ‘crowd’ around the resident AZ or supplemented nearby secondary AZs. The latter predominantly constituted directly neighboring synapses, but also included distal synapses that were spatially separated by significant distances (∼7 µm). Even in such cases, the transfer appeared highly directional and targeted. Importantly, intersynaptically exchanged precursors contained the CAZ proteins Piccolino and Bassoon in addition to RIBEYE, and thus would rapidly modulate the available SV contingent. While exchange of presynaptic proteins and SVs between neighboring boutons has previously been described in neurons – where it gives rise to continuous fluctuations in individual synapse composition ^55–58^ – such re-distribution events occurred with much slower pace than the ones observed here (e.g., >8 h for Bassoon ^56^). Hence, the assembly into precursor organelles may present a unique and highly efficient mechanism for rapid structural adaptation.

### Putative mechanisms of ribbon scaffold assembly

The finding that even the smallest precursors appeared to constitute multi-protein organelles questions how they initially assemble and structurally reorganize during plasticity events. While it is well established that RIBEYE auto-aggregates in the cytosol of sensory receptor cells ^11,13,18,39^, it has remained unclear how the other CAZ proteins are recruited to nascent ribbons. RIBEYE aggregates also form in heterologous expression systems ^59^, and a recent study could show that – once supplemented with other CAZ components including Bassoon, RIMBP2 and CaV1.3 – artificial membrane-anchored ribbons could be generated autonomously ^60^. Here, the self-organizing properties of IHC CAZ proteins are reminiscent of nanoclusters/nanocolumn formation at central synapses, where the association of different proteins in biomolecular condensates has been suggested to play a significant role in AZ assembly via liquid-liquid phase separation (LLPS ^61^). Remarkably, ribbon precursors meet multiple criteria regarding LLPS: (i) precursors are highly dynamic and exhibit a tendency for self-association and spontaneous reversible dispersion, (ii) they occur in mobile and immobile fractions that can exchange material, (iii) ribbons display scaffold-intrinsic mobility, (iv) RIBEYE contains disordered protein sequences and Proline-rich regions, that likely contribute to (v) the ability of RIBEYE to interact with other RIBEYE molecules through various binding sites ^32,59,62,63^. Moreover, similar to our previous findings of ribbon precursor translocation ^14,41^, other biomolecular condensates – such as ribonucleoprotein granules – have been shown to be transported along MTs ^64^. This is an intriguing combination of intracellular displacement mechanisms, which could explain the highly variable mobility patterns we detected for ribbon precursors during development. Finally, an interesting feature proposed for LLPS taking place at conventional AZs is the implication of two functional pathways: one for the clustering of CaVs and one for SV organization ^65^. The unique position of the synaptic ribbon might bridge these functionalities by clustering CaV1.3 and facilitating SV priming and replenishment ^4,8,10^. Altogether, this makes for an enticing argument for ribbon precursors as biomolecular condensates, and a possible contribution of LLPS to IHC synapse assembly. Future studies will be needed to test this hypothesis.

### Activity-dependent plasticity of developing ribbon synapses

Finally, we investigated a functional link between ribbon plasticity and spontaneously-occurring pre-sensory SA. Considering the strong interconnectivity between anchored ribbons, CAZ proteins and CaV clusters, it is conceivable that activity-dependent Ca^2+^ influx and/or the associated membrane turnover could alter ribbon dynamics. In fact, previous studies reported a correlation between synaptic activity and CaV cluster mobility along the presynaptic membrane of retinal photoreceptors ^66^. In IHCs, pharmacological silencing of SA significantly attenuated precursor plasticity at the AZ, as synapse-interacting tracks displayed plasticity profiles comparable to non-synaptic precursors, thus implicating CaV1.3-mediated Ca^2+^ influx as a major driver of this process. Interestingly, various molecular motors have been shown to be regulated by local calcium levels, to either associate or disassociate from the cargo or the cytoskeleton ^40^. Likewise, the formation of distinct liquid states during LLPS can depend on local increases in Ca^2+^ levels ^67^, thus rendering both mechanisms as plausible candidates for the observed bi-directional plasticity of ribbon scaffolds.

Interestingly, at conventional synapses, synaptic activity has been shown to exert differential effects on the presynaptic protein complement: while Munc-18 and Synapsin were redistributed between synapses in an activity-dependent manner ^57,68^, Munc-13 and Bassoon remained unaffected – thus indicating an mechanistic distinction between soluble SV-associated proteins and membrane-anchored scaffolds. This places the ribbon in a unique position, being involved in both pathways. However, pharmacological silencing did not only affect ribbon precursors in close proximity to the AZ but indiscriminately reduced precursor velocity and limited their volume acquisition. The disruption of ribbon precursor dynamics regardless of their synaptic engagement suggests that this effect is separate from the AZ-centered attenuation of structural plasticity, and likely involves a fundamental molecular pathway of IHC homeostasis. Here, an interesting comparison is the genetic or pharmacological inhibition of presynaptic Ca^2+^ influx in zebrafish nHCs, which lead to an increase in ribbon precursor volume in young nHCs ^36,69^. While opposite in directionality, the impact of isradipine illustrates a seemingly conserved mechanism between presynaptic Ca^2+^ influx and structural plasticity. Finally, the apparent correlation between presynaptic Ca^2+^ levels and precursor plasticity is of interest in the context of the location-dependent plasticity we detected along the pillar/modiolar IHC axis: The highly plastic and more ‘synaptic-like’ characteristics of ribbon precursors in the modiolar IHC fraction imply a potentially higher level of Ca^2+^ influx driving this difference. This would align with the higher number and volume of modiolar ribbon synapses found in mature IHCs, which have been shown to correlate with the amplitude of Ca^2+^ influx, number of CaVs, and their activation thresholds ^33,50,51,70–72^. However, it remains unclear whether any heterogeneous CaV expression pattern is already present during early development and if distinct innervation may retrogradely initiate the differentiation of modiolar AZs. Yet, the exact driver of such reciprocal pre- and postsynaptic influence remains to be determined.

Together, our data substantiate previous work indicating intrinsic synaptic differences along the pillar/modiolar axis during early development ^34^ and this pre-determined polarity appears fundamental for adequate AZ development.

### Technical limitations of this study

The cochlea is tightly embedded in the temporal bone and hence inaccessible for high-resolution long-term optical analysis of individual ribbon synapses *in vivo*. Thus, we opted for a short-term organotypic culture approach to study the mechanisms of synaptogenesis in a tightly-controlled semi-intact system. While our organotypic cultures retained excellent structural integrity and afferent innervation, the dissection and culturing procedure nevertheless causes significant loss of SGNs during the immediate culturing period. Yet, through limiting our analyses to a brief and defined time window subsequent to tissue extraction and carefully monitoring cell health and global structural morphology during image acquisition, we ensured tissue preservation in a near-native state. While these precautions enabled us to conduct detailed analyses of developing IHCs *in situ*, we faced several methodological limitations that need to be considered: Firstly, the temporal resolution of multi-color 3D data acquisition is severely restricted by the sequential fluorescence channel acquisition. Therefore, ultrafast events will have failed to register in our experiments and rapid displacements or transient plasticity events may have been missed. Future studies should hence opt for high-speed imaging techniques to extend our knowledge on nanoscopic protein dynamics during IHC synaptogenesis. Secondly, the Halo-based PSD labeling was variable between experiments and differed in SNR (i.e., the JF549 ligand consistently outperformed the JF646 ligand in our triple-color experiments). Hence, we cannot exclude the possibility that dimly fluorescent PSDs went undetected or even misclassified as ‘membrane-proximal’ instead of ‘synaptically-engaged’. Although this may minimally alter the population statistics of membrane-proximal precursors, it would not affect the synaptic class that partner a bright, distinguished PSD.

## Acknowledgements

We would like to express our utmost gratitude to Christiane Senger-Freitag and Sandra Gerke for expert technical support – in particular CSF for performing the intracochlear injections – and Kathrin Kusch for additional AAV production. Also, we would like to thank the City Campus Light Microscopy Facility at the Max Planck Institute for Multidisciplinary Sciences for making our live-cell imaging experiments possible. pUCmini-iCAP-PHP.eB was a gift from Viviana Gradinaru (Addgene plasmid # 103005; http://n2t.net/addgene:103005; RRID:Addgene_103005). This work was funded by the Collaborative Research Center 889 “*Cellular Mechanisms of Sensory Processing*” from the Deutsche Forschungsgemeinschaft (DFG, German Research Foundation): project B08 to CV and an Otto Creutzfeldt Fellowship of the Elisabeth and Helmuth Uhl Foundation to CV. Part of this research was funded by the Austrian Science Fund (FWF) 10.55776/PAT2555923 to CV.

Mouse line and HaloTag work was supported by funding from the European Research Council (ERC) under the European Union’s Horizon 2020 Research and Innovation Programme (695568 SYNNOVATE; 885069 SYNAPTOME) and a Wellcome Technology Development grant (202932) to S.G.N.G.; and by the Simons Foundation Autism Research Initiative (529085) to N.H.K. and S.G.N.G. For the purpose of open access, the author has applied a CC-BY public copyright license to any Author Accepted Manuscript version arising from this submission.

Work in the Wolf lab is supported by the Deutsche Forschungsgemeinschaft (DFG, German Research Foundation) 436260547 in relation to NeuroNex (National Science Foundation 2015276) & under Germany’s Excellence Strategy - EXC 2067/1- 390729940, by the DFG - Project-ID 317475864 - SFB 1286, Project-ID 454648639 - SFB 1528, Project-ID 273725443 - SPP 1782, and Project-ID 430156276 - SPP 2205. This work was further supported by the Leibniz Association (project K265/2019), and by the Niedersächsisches Vorab of the VolkswagenStiftung through ZN3420 and through the Göttingen Campus Institute for Dynamics of Biological Networks (ZN3326 & ZN3371).

## Author contributions

C.V. conceived the project, C.V. and R.A.V. designed the experiments. R.A.V. performed live-cell imaging, immunohistochemistry, confocal/STED imaging and data analysis. M.S., J.B. and F.W. performed additional data analysis. N.H.K. and S.G.N.G. contributed mouse lines and HaloTag labelling support. V.R. designed viral constructs and generated AAVs. R.A.V. and C.V. wrote the paper and generated all figures. All co-authors revised the manuscript.

