## Supplemental Information for "Developmental plasticity facilitates the structural maturation of cochlear inner hair cell ribbon synapses"

### Supplemental Tables:

**Table ST1. Primary, secondary and conjugated antibodies used in experimental procedures.**

| Primary ABs | Species | Brand | Cat. Numb. |
|---|---|---|---|
| αβ-3-Tubulin (Tuj-1) | Mouse IgG2a | BioLegend | #801202 |
| αBassoon | Chicken | Synaptic Systems | #141016 |
| αBassoon | Mouse IgG2a | Abcam | #ab82958 |
| αCtBP2 | Mouse IgG1 | Becton Dickinson | #612044 |
| αMyosin VIIa | Mouse IgG1 | DSHB | #MYO7A 138-1 |
| αPiccolino (αAczp18p19) | Rabbit | a kind gift of Prof. Manfred Kilimann [1,2] | Limbach et al. 2011 |
| αPSD95, clone 7E3-1B8 | Mouse IgG1 | Sigma | #MAB1598 |
| αRIBEYE-A | Rabbit | Synaptic Systems | #192103 |
| **Secondary ABs** | **Species** | **Brand** | **Cat. Numb.** |
| αChicken AlexaFluor647 | Goat | Abcam | #ab150171 |
| αMouse Star580 | Goat | Abberior | #ST580-1001-500UG |
| αMouse IgG1 AlexaFluor488 | Goat | Life Technologies | #A21121 |
| αMouse IgG1 AlexaFluor594 | Goat | Life Technologies | #A21125 |
| αMouse IgG1 Star635P FluoTag®-X2 | Camelid | NanoTag, | #N2002-Ab635P |
| αMouse IgG2 Atto565 FluoTag®-X2 | Camelid | NanoTag, | #N2702-At565 |
| αMouse IgG2 Star635P FluoTag®-X2 | Camelid | NanoTag | #N2702-Ab635P |
| αRabbit Star580 FluoTag®-X4 | Camelid | NanoTag | #N2404 |
| αRabbit Star635P | Goat | Abberior | #ST635P-1002-500UG |
| **Conjugated ABs** | **Species** | **Brand** | **Cat. Numb.** |
| αGFP Atto488 FluoTag®-X4 | Camelid | NanoTag | #N0304-At488 |
| αPSD95 Atto488 FluoTag®-X2 | Camelid | NanoTag | #N3702-At488 |
| αPSD95 AlexaFluor647 FluoTag®-X2 | Camelid | NanoTag | #N3702-AF647 |
| αRFP Atto565 FluoTag®-X4 | Camelid | NanoTag | #N0404-At565 |

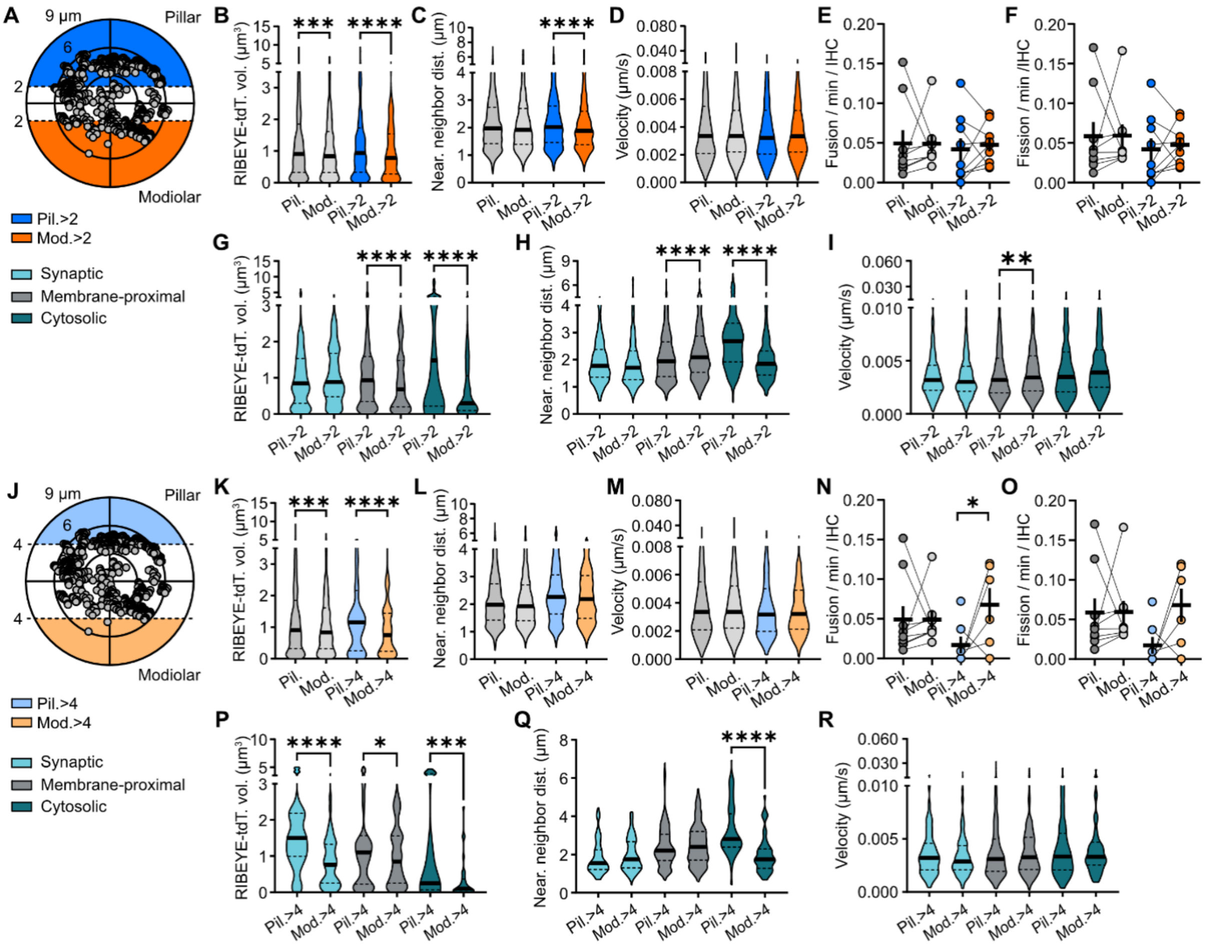


**Figure S1. Ribbon plasticity along the pillar/modiolar IHC axis, cutoffs 2 µm and 4 µm. A.** Illustration of the >2 µm extremities within the axes of the IHC; all precursors over time are displayed. **B-F.** Grays: pillar/modiolar division based on the 0 midline, blue/orange: pillar/modiolar populations taken from the >2 µm extremities. **B.** The volume of modiolar ribbon precursors is lower than of pillar precursors. **C.** The nearest neighbor distance of modiolar precursors is lower than of pillar precursors in the >2 µm extremities. **D.** Ribbon precursor velocity is comparable throughout the IHC. **E, F.** The frequency of fusion (F) and fission (G) events per minute per IHC is comparable in modiolar and pillar tracks in the >2 µm IHC extremities. **G-I.** Specifically for the >2 µm extremities, considering the three different precursor classes: synaptic (cyan), membrane-proximal (gray), and cytosolic (teal) precursors. **G.** The volume of membrane-proximal and cytosolic modiolar precursors was lower than of pillar precursors. **H.** The lower nearest neighbor distance of the >2 µm modiolar precursor population appears to be driven by the cytosolic precursors, whereas the membrane-proximal precursors are in fact at a slightly higher nearest neighbor distance. **I.** The ribbon precursor velocity was slightly higher for modiolar membrane-proximal precursors than pillar, and showed a similar trend for the cytosolic precursor population ($p=0.0518$), in the >2 µm pillar/modiolar IHC extremities. **J.** Illustration of the >4 µm extremities within the axes of the IHC; all precursors over time are displayed. **K-O.** Grays: pillar/modiolar division based on the 0 midline, light blue/orange: pillar/modiolar populations taken from the >4 µm extremities. **K.** The volume of modiolar ribbon precursors is lower than of pillar precursors. **L.** The nearest neighbor distance of modiolar precursors does not differ from pillar precursors in the >4 µm extremities. **M.** Ribbon precursor velocity is comparable throughout the IHC. **N.** The frequency of fusion (F) events is higher in the modiolar >4 µm IHC extremities than the pillar. **O.** The number of fission (G) events per minute per IHC shows a trend towards a higher frequency in the modiolar than pillar tracks in the >4 µm IHC extremities ($p=0.0641$). **P-R.** Specifically for the >4 µm extremities, considering the three different precursor classes. **G.** The volume of membrane-proximal and cytosolic modiolar precursors was lower than of pillar precursors. **H.** The nearest neighbor distance of the >4 µm modiolar precursor only differs from the pillar population in the cytosolic precursor class. **R.** The ribbon precursor velocity was similar for all precursor classes in the >4 µm pillar/modiolar IHC extremities. *$p<0.05$, **$p<0.01$, ***$p<0.001$, ****$p<0.0001$. Statistical significance: Kruskal-Wallis. $N_{animals}=4$, $n_{IHC}=9$, $n_{ribbons}=10855$.

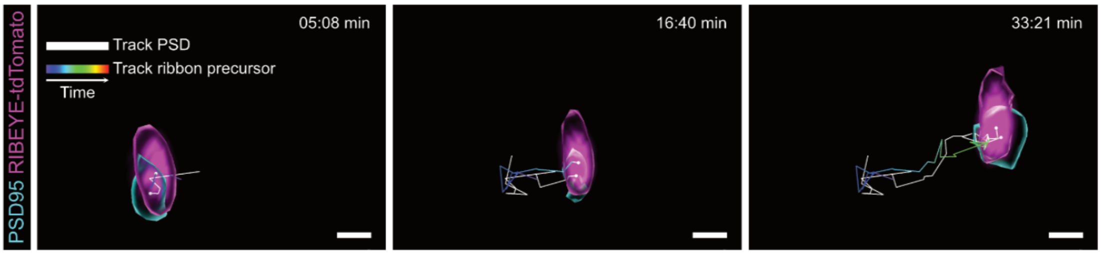


**Figure S2. Co-mobility of a trans-synaptically coupled ribbon precursors and PSD.** Exemplary 3D live imaging frames of a ribbon precursor (magenta) engaged at the presynaptic AZ and PSD cluster (cyan), traced during coupled displacement. The aligned tracks are shown in white (PSD) and a rainbow color gradient (ribbon precursor), reflecting the time. Scale bar: 1 µm.

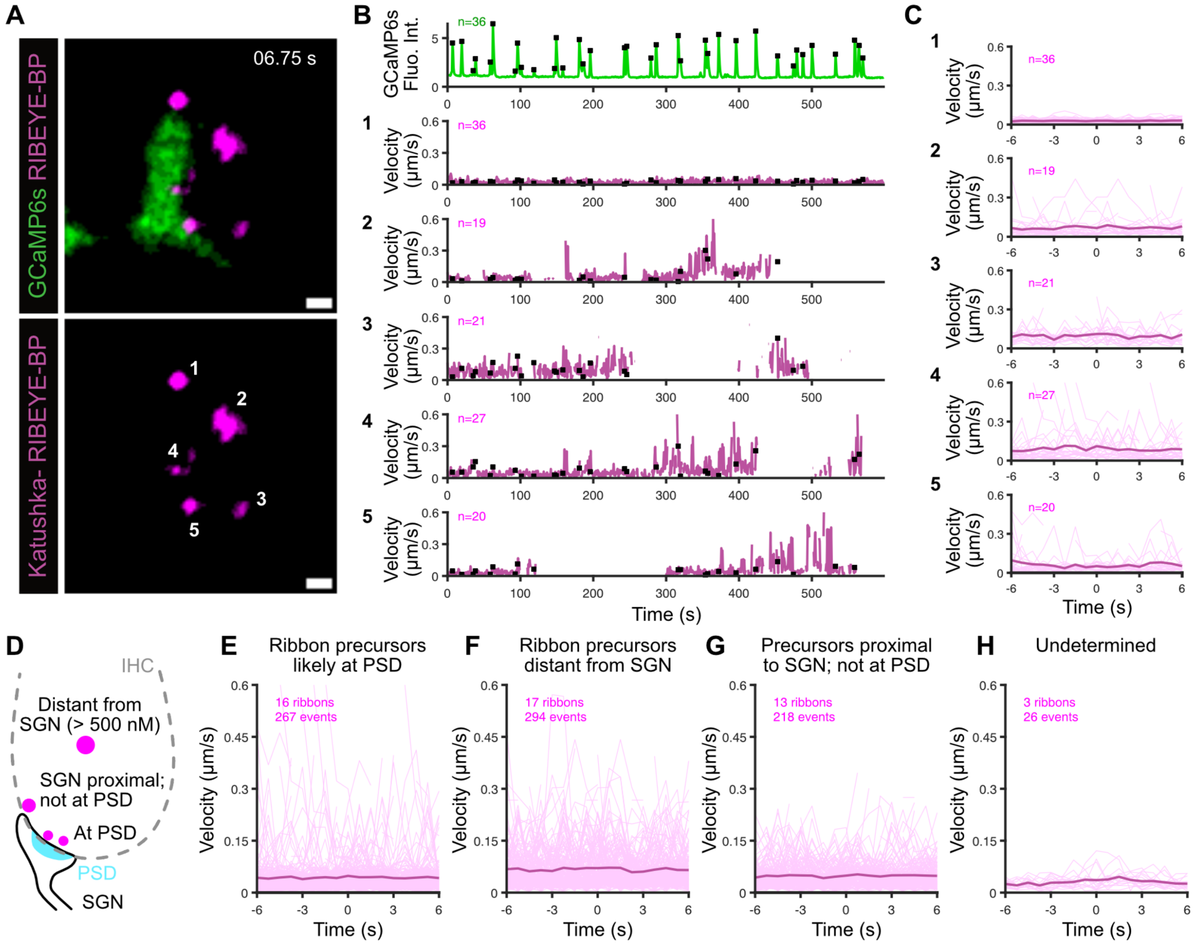


**Figure S3. Ribbon precursor dynamics do not correlate with postsynaptic SGN activity. A.** Exemplary live imaged frame of an IHC expressing Katushka-RIBEYE-binding peptide (P3DIV3), innervated by a GCaMP6s expressing afferent SGN. The different traced ribbon precursors are indicated by number. **B.** GCaMP6s fluorescence intensity was traced over time as a postsynaptic measure of spontaneous synaptic activity and exocytotic release of the IHC. Simultaneously, the dynamics of the ribbon precursors indicated in A were traced, plotted here is the object velocity over time. Interludes of trace absence likely reflect the displacement in Z of the ribbon precursor, out of the imaged ROI (single plane). **C.** Spike-triggered averaging of velocity traces using the GCaMPs fluorescence as trigger (12 s time window). The number of events used for averaging is indicated by 'n', reflecting the number of fluorescence peaks that could be correlated to ribbon precursor velocity values. **D.** Schematic illustration of different ribbon precursor categories based on their intracellular location. **E-H.** Grand averages of spike-triggered velocity traces for ribbon precursors of the different categories in D: precursors (**E**) localized in probable association with a PSD; (**F**) at a distance (> 500 nM) from the fluorescent afferent SGN; (**G**) proximal to the innervating SGN, but localized where partnering with a PSD is highly unlikely; (**H**) of undetermined location. The absence of a reproducible pattern suggests statistical independence between fluorescence dynamics and ribbon mobility, in all precursor locations. $N_{animals}$= 3, $N_{IHC}$= 8, $n_{ribbons}$= 49. Scale bars: 1 µm.

**Methods Figure S3. Time-lapse of ribbon dynamics combined with SGN $Ca^{2+}$ imaging**

**Viral transduction**

Viral transduction *in vitro* was performed on P3 or P4/DIV0 cultures of SNAP25GCaMP6s mice, by application of PHP.B_HBA_Katushka-RIBEYE-BP_WPRE_bGHpA (1.5e+11 gc/mL) and incubation at 37C 5%CO2 for ≥72 hours (DIV3).

**Short-term high-speed dual-color time-lapse imaging**

Short-term time-lapse imaging experiments were performed at an Abberior Instruments Expert Line STED microscope, operated in confocal laser scanning mode, using a 60x/1.20 NA water immersion objective. A top mount on-stage incubator (Okolab uno stage top incubator, H391-Olympus-IX-SUSP 2015) was used to create and maintain environmentally-controlled conditions (37°C, 5% CO2). ROI selection was based on GCaMP6 signal strength and fluorescence periodicity, plus, when present, detection of the Katushka-RIBEYE-BP expression levels to be sufficient for imaging. Dual-color time-lapse images were acquired over a 10-minute period, and consisted of single-plane optical sections, at intervals of 0.75 seconds. Spatial dimensions of the regions for either imaging condition were minimized and adapted to tissue orientation, at approximately 15x15 µm. The two colors were acquired in confocal mode, by sequential line scanning, 2 times accumulation and a 5 ns dwell time.

**Time-lapse image processing and analysis**

The fluorescent GCaMP6s signal was processed in FIJI/ImageJ, by extraction of the fluorescence intensity (F) over time. To create the ROI, an image mask was generated based on the maximum intensity projection, followed by intensity thresholding and noise reduction. The extracted fluorescence intensity was then baseline corrected ($\Delta F = F - F_0$). The ribbon precursor velocity was traced in IMARIS, using the Spots function. Potential synchronicity between peaks in the GCaMP6s fluorescence and ribbon precursor velocity was analyzed using custom MATLAB scripts.
